# Survival curve steepening and flattening reflect evolutionarily conserved changes in the variability of aging

**DOI:** 10.64898/2026.02.15.705972

**Authors:** Bruce Zhang, Gillian Borland, Yiran Zhang, Zibo Gong, Colin Selman, David Gems

## Abstract

As global human life expectancy continues to rise, accompanying increases in healthspan that prevent morbidity expansion become increasingly imperative. Population lifespan can increase in distinct ways, for instance through rectangularization (steepening) or triangularization (flattening) of survival curves. These two demographic changes, particularly rectangularization, occur frequently across human and model organism populations, yet their biological determinants and effects on healthspan and morbidity are largely unknown. Notably, these modes of life-extension occur when parameters of the Gompertz mortality model (capturing exponential age-increases in mortality rate) change inversely, a widely-reported phenomenon known as the Strehler-Mildvan correlation – whose biological basis also remains unexplained. We therefore investigated longitudinal health, morbidity and lifespan in 30 *Caenorhabditis elegans* cohorts using multiple life-extension protocols. We report that survival curve rectangularization results from healthspan expansion in short-lived population members, whereas triangularization (which primarily extends the curve tail) from healthspan and morbidity expansion in long-lived population members. Interestingly, rectangularization and triangularization respectively decrease and increase inter-individual variation in the aging process, and the mode of life-extension that occurs depends on levels of existing variation. Notably, triangularization was more effective at extending lifespan without morbidity expansion. Analysis of fruit fly and mouse data show that these biodemographic dynamics are also largely evolutionarily conserved.

## Introduction

The biological process of aging (senescence) is a leading cause of mortality and morbidity worldwide (1), yet its causes and mechanisms remain poorly understood. Given continued increases in global life expectancy (2), concomitant changes in the durations of health (healthspan) and morbidity, and their biological determinants, become increasingly important to quantify and understand.

Lifespan increases can be studied through mortality modelling, such as with the Gompertz equation (3), *μ(x)=αe^βx^*, which can capture the exponential increase in mortality rate observed in many animal species, including humans. Here, mortality rate at age *x* is determined by a scale parameter *α* and rate parameter *β*, which respectively specify the theoretical initial mortality rate (at *x*=0) and mortality rate acceleration with advancing age. These parameters have different effects on the survival curve (Fig. 1A). Although the Gompertz model is frequently employed in aging research, the biological meaning of *α* and *β* has only recently been demonstrated empirically (4).

**Fig. 1.**
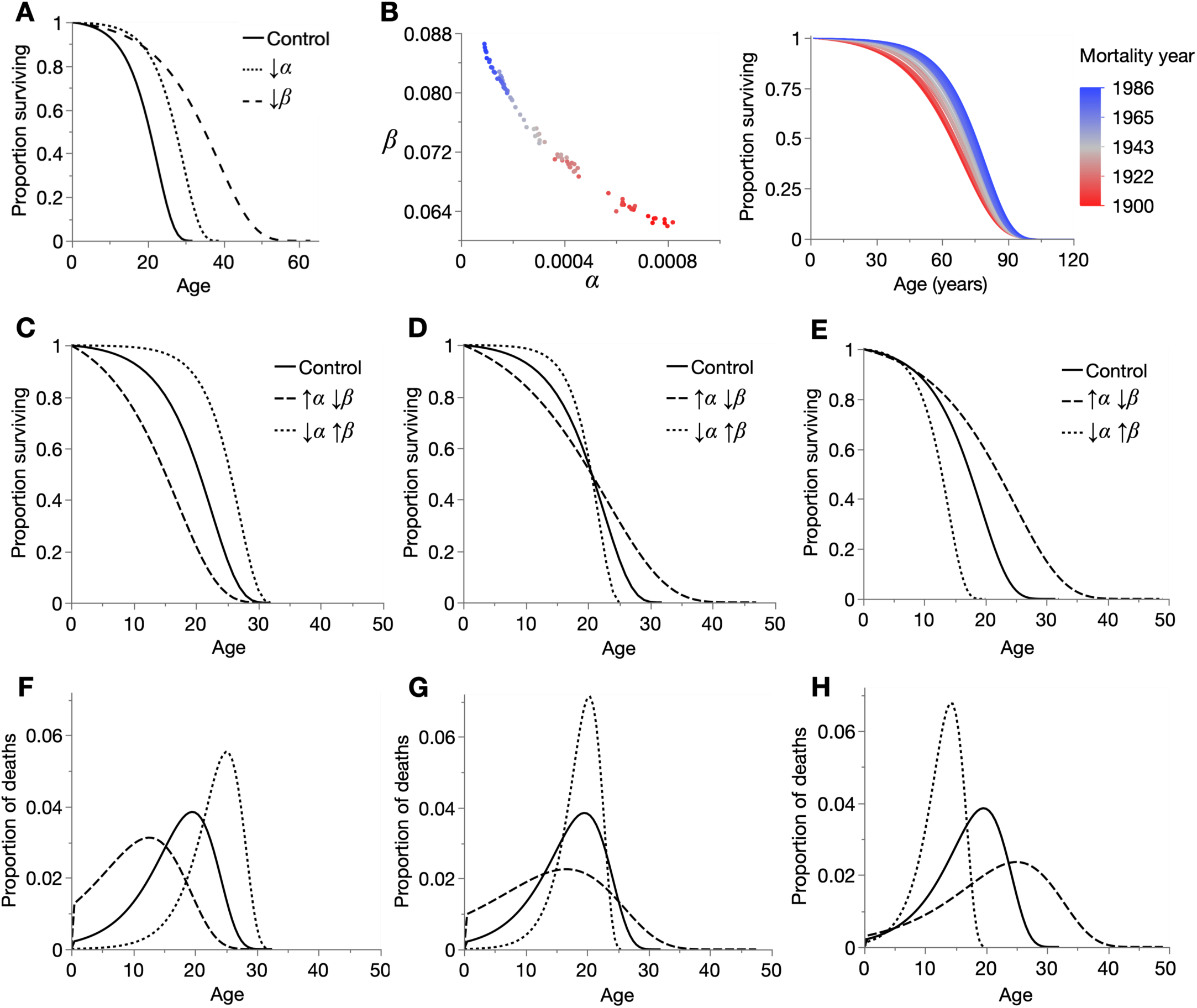
Demographic effects of inverse changes between Gompertz parameters. (**A**) Effect of decreasing Gompertz parameters *α* and *β* on the survival curve. **Control**: *α*, *β*: 0.002, 0.2; **↓*α***: *α*, *β*: 0.0005, 0.2; **↓*β***: *α*, *β*: 0.002, 0.1. (**B**) Left: inverse relationship between the Gompertz parameters, forming a Strehler-Mildvan (S-M) correlation, from annual period mortality (i.e. all deaths that year). Right: equivalent Gompertz survival curves, of U.S. men between 1900 and 1986, produced from data in Riggs (6). (**C**–**E**) Three forms of survival curve transformation that occur in S-M correlations: (**C**) rectangularization/de-rectangularization, (**D**) intersection of survival curves, and (**E**) triangularization/de-triangularization. In these examples, *α* and *β* have been set so that maximum (**C**), median (**D**) and minimum (**E**) lifespans are approximately equivalent, but these values can vary. The parameters are: (**C**) **Control**: *α*=0.002, *β*=0.2; **↑***α***↓***β*: *α*=0.012, *β*=0.14; ↓*α***↑***β*: *α*=0.0001, *β*=0.29, (**D**) **Control**: *α*=0.002, *β*=0.2; **↑***α***↓***β*: *α*=0.012, *β*=0.08; **↓***α***↑***β*: *α*=0.0001, *β*=0.37, and (**E**) **Control**: *α*=0.002, *β*=0.2; **↑***α***↓***β*: *α*=0.003, *β*=0.12; **↓***α***↑***β*: *α*=0.0012, *β*=0.35. (**F**–**H**) Respectively, effects of Gompertz parameter changes in **C**–**E** on mortality frequency over time.

A further enigma is the Strehler-Mildvan (S-M) correlation: an inverse correlation between *α* and *β*, that is frequently observed between human populations (5–7), most notably in the steepening of global survival curves over the last two centuries (Fig. 1B). The correlation has since also been reported between laboratory animal populations (8–11). In a modified form, the S-M correlation is also known as the compensation law of mortality (12).

The S-M correlation has distinctive effects upon the survival curve, for instance causing rectangularization/de-rectangularization (Fig. 1C, 1B right), intersection (crossing over) (Fig. 1D), or triangularization/de-triangularization (Fig. 1E). The latter term (triangularization) we introduce in this study, to describe the selective extension of the survival curve tail, which produces a flattened curve resembling the protruding corner of a triangle. Lifespan extension can therefore occur through decreased *α* and increased *β* (rectangularization) or through increased *α* and decreased *β* (triangularization); in the life-shortening direction, these changes become de-rectangularization and de-triangularization, respectively. Importantly, because the Gompertz parameters affect lifespan in the same direction (decreasing *α* or *β* both increase lifespan; Fig. 1A), S-M correlations always involve antagonistic lifespan effects. Thus, depending on the magnitudes of inverse *α* and *β* change, survival curves can converge anywhere along their lengths: at late ages in rectangularization (Fig. 1C), early ages in triangularization (Fig. 1E), or somewhere in between in the case of intersection (Fig. 1D). These S-M correlations have intuitive effects upon the mortality frequency distribution, which is approximately shifted and stretched/compressed by *α* and *β*, respectively (Fig. 1F–H).

However, the biological basis of the S-M correlation, in terms of changes in the aging process, remains unclear and has not been determined empirically. Possible explanations include the effects of antagonistic pleiotropy (13) and intra-population heterogeneity (13–15). S-M correlations can also arise from modeling artifacts (16–18) but, as we will show, not in the data we present here. Importantly, effects of rectangularization and triangularization, the demographic building blocks of S-M correlations, on longitudinal durations of healthspan and morbidity have not been directly studied.

We recently investigated the biological basis of the Gompertz parameters in 24 differently-lived cohorts of *C. elegans*. Our findings showed that, contrary to long-standing theoretical interpretations, *α* and *β* respectively reflect biological aging rate and inter-individual variability in morbidity duration (4). In the present study, we revisit and add to these earlier cohorts to examine empirically the biological changes underpinning the survival curve transformations that occur in S-M correlations. We find that survival curve rectangularization reflects healthspan expansion in shorter-lived population members and increased homogeneity in the aging process. In contrast, survival curve triangularization reflects both healthspan and gerospan (morbidity) expansion in longer-lived population members and increased heterogeneity in the aging process. Notably, similar biodemographic dynamics underly these survival curve transformations in fruit fly and mouse populations. Our findings present an empirical, biological explanation of survival curve steepening and flattening (see Fig. 9 for a schematic overview), whose determinants may be conserved between invertebrates and mammals, and potentially underly the Strehler-Mildvan correlation.

## Results

### Lifespan extension by rectangularization and triangularization in *C. elegans*

Despite frequent use of *C. elegans* in Gompertzian mortality modeling, their utility for understanding the S-M correlation remains largely unexplored. To address this, we produced a longitudinal health and mortality dataset of 30 nematode cohorts, comprised of all combinations of 3 culture temperatures (15°C, 20°C, 25°C), with/without antibiotic (carbenicillin), and 5 genotypes (wild type, *daf-2(m577)*, *daf-2(e1368)*, *daf-2(e1370)*, *daf-16(mgDf50)*). We recently used 24 of these cohorts to investigate the biological basis of the Gompertz parameters (4), and here present the completed dataset that also includes 6 *daf-16* cohorts (at 3 temperatures, with/without carbenicillin), additional necropsy data (of end-of-life pathology), and measurements of early-adulthood bacterial lawn (food) interactions for each cohort.

Gompertz parameters were obtained from the lifespan data of each cohort by maximum likelihood estimation (19) (Table S1). These Gompertz fits were verified with parametric Anderson-Darling goodness-of-fit tests (Table S2), and additionally predicted first quartile, median and third quartile lifespans with R^2^ of 0.981, 0.996, 0.993, respectively (Fig. S1). Each cohort (mean *n*=144 individuals/cohort) was pooled from at least 3 sequential trials, whose inter-trial lifespan variation was low (Fig. S2, Table S1). We also previously showed that increasing population sizes for 24 of these cohorts by ∼3-fold (>300 individuals/cohort) had negligible effects on lifespan and Gompertz parameter estimates (4).

The 30 cohorts yield in total 435 pairwise comparisons between them, each reflecting a distinct biological change in the aging process and lifespan. We therefore refer to these 435 comparisons as *treatments* and evaluate them in the lifespan-extending direction (i.e. 435 lifespan-extending treatments). This pairwise approach not only informs about lifespan extension by specific conditions/mechanisms (temperature, antibiotics, IIS pathway mutations, and their interactions) but makes possible the study of condition-independent patterns, i.e. fundamental principles that govern the relationship between demographic and biological aging.

We began by assessing the effects of these 435 lifespan-extending treatments on the Gompertz parameters, *α* and *β* (Fig. 2A). 172/435 (40%) of treatments increased lifespan by decreasing both Gompertz parameters (except for 3 treatments that had no effect on *α*), while the majority, 263/435 (60%), increased lifespan through inverse parameter changes: 101 treatments decreased *α* and increased *β* (i.e. rectangularization), and 162 increased *α* and decreased *β* (i.e. triangularization). Thus, lifespan extension amongst these 30 cohorts arises predominantly through survival curve transformations that occur in S-M correlations: rectangularization which steepens the survival curve (Fig. 1C), and triangularization which flattens it (Fig. 1E). We will refer to these as rectangularizing and triangularizing treatments, and the remaining 172 treatments (in which the Gompertz parameters did not vary inversely) as non-S-M treatments.

**Fig. 2.**
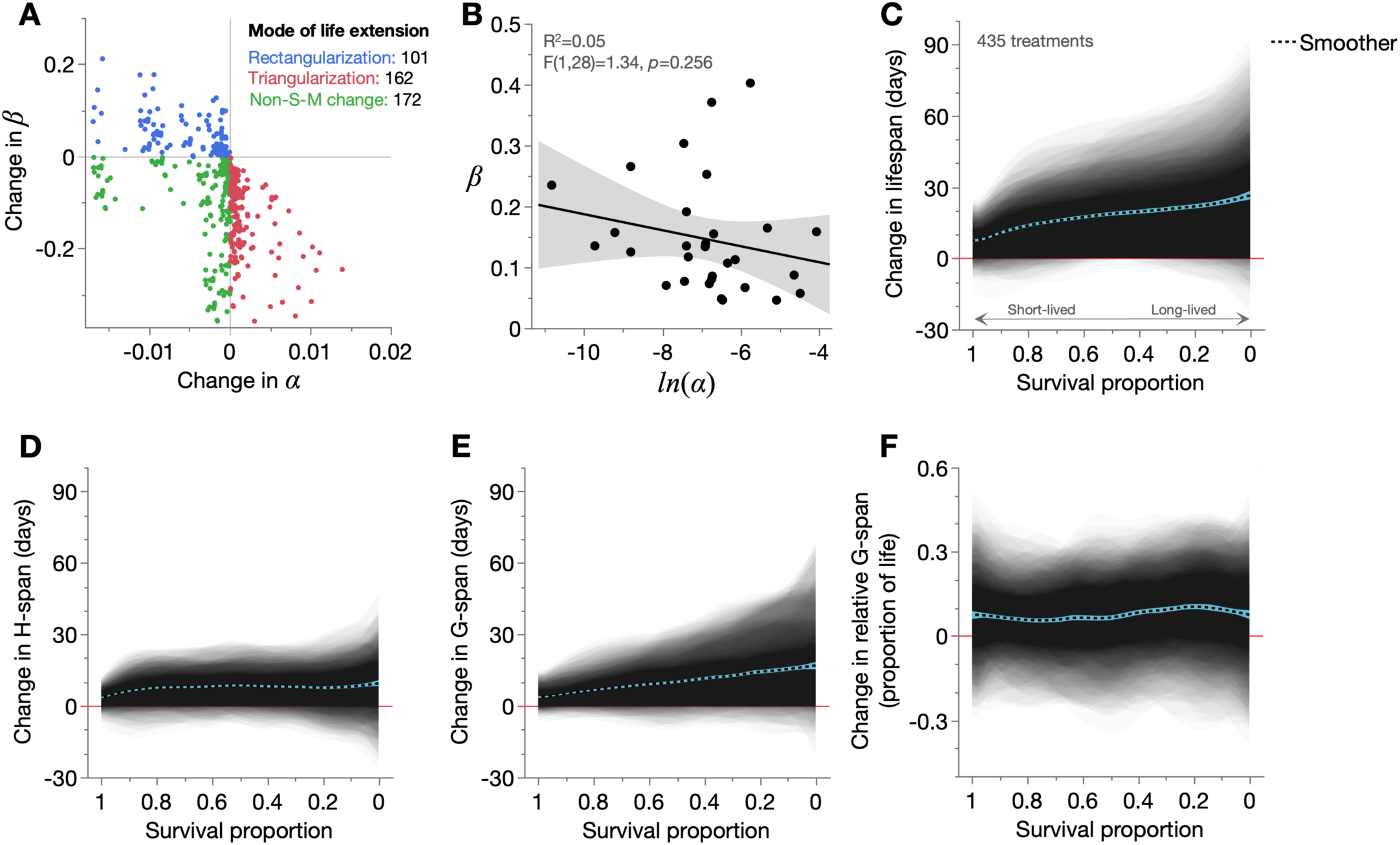
Demographic modes of lifespan extension and their effects on healthspan and gerospan. (**A**) Changes in *β* between all possible pairs of the 30 cohorts, over the corresponding change in *α* for those pairs. Change direction was always determined based on the longer-lived cohort minus shorter-lived cohort, such that all changes reflect the effects of lifespan extension. (**B**) Least-squares linear regression of the relationship between *α* (ln-transformed) and *β* across the 30 cohorts, with the 95% confidence region shaded and F-test statistics annotated. (**C**–**F**) Shaded and overlaid area plots for all 435 lifespan-extending treatments (comparison pairs), of changes in (**C**) lifespan, (**D**) H-span, (**E**) G-span, and (**F**) relative G-span. Each of the 435 pairwise comparisons contributes one shaded change (JMP transparency=0.02), which is the area between the lifespan (or H-span, G-span, relative G-span) distributions of the control and treatment cohorts in that pairwise comparison. These shaded changes are plotted over survival proportion (x-axis left: shorter-lived population members, x-axis right: longer-lived population members), enabling visualization of in which population members a particular change (to lifespan, H-span, G-span, or relative G-span) occurs. Red line: y=0, below which the lifespan-extending treatment shortens that trait. Dashed black line: summary spline smoother (lambda=0.05) of the 435 treatment changes, showing its 95% confidence region in blue.

As expected, no treatments increased both Gompertz parameters (which would decrease overall lifespan) (Fig. 2A), reaffirming that the estimated parameters accurately capture lifespan changes in these treatments. This, and the robust fitting of the Gompertz model to these cohorts (Fig. S1, Table S2) show that the rectangularization and triangularization occurring in this dataset are not statistical modeling artifacts, which may occur under certain conditions (16–18). Here, our treatments reflect real (biological) changes in lifespan and thus also the aging process. Furthermore, exclusion of treatments in which one or both Gompertz parameters were not statistically significantly changed predominantly excluded non-S-M treatments (Fig. S3).

We also checked whether an overall S-M correlation was observed across the 30 cohorts. We did not observe a statistically significant inverse relationship between the Gompertz parameters (Fig. 2B), consistent with the presence of treatments (40%) that decreased both parameters. Therefore, the dataset, generated from combinations of diverse interventions (temperature, antibiotics and nutrient signaling), presents a mixture of demographic modes of life-extension and an opportunity to understand their differences. This suggested that our investigation could potentially reveal fundamental biological principles underlying rectangularization and triangularization, including those occurring in S-M correlations.

### Different dynamics of health and morbidity during lifespan extension

We then addressed the focus of this study: the biological features of the aging process that occur in rectangularization and triangularization. To this end, alongside collecting lifespan data, we longitudinally tracked the health of each individual nematode from reproductive maturity until death, in all 30 cohorts. This included assessment of locomotory capacity every 2–3 days, interaction with the *E. coli* food source every 2–3 days, and *E. coli* infection status at the time of death. Such simultaneous quantification of biological aging traits and lifespan in the same individuals can reveal the biological basis of demographic mortality patterns (4, 20).

We first visualised the effects of the 435 treatments on lifespan, by shading the change in survival curves for each treatment, and overlaying them, plotted over a survival proportion x-axis that ranks population members in order of increasing lifespan, from left to right (Fig. 2C). This mode of presentation shows in which population members a change of interest (here, lifespan) occurs. As expected, the 435 treatments increased lifespan and, as often observed in nematode survival studies (4, 21, 22), more so in already longer-lived population members (Fig. 2C). Lifespan of the shortest and longest-lived population members was also modestly decreased in some treatments, indicating antagonistic lifespan effects (i.e. intersecting survival curves).

We similarly examined effects of these treatments on aging-related health and morbidity, here respectively measured as youthful (sinusoidal) and reduced (non-sinusoidal or immotile) locomotory capacity, which are physiologically-integrative measures of key stages in the aging process (4, 23, 24). We refer to the number of days of life spent in youthful and reduced locomotory capacity as *healthspan* (H-span) and *gerospan* (G-span), respectively, and the proportion of life spent in gerospan as *relative gerospan* (relative G-span). Because H-span + G-span = lifespan, we can explain changes in lifespan in terms of changes in H-span and G-span (i.e. a biological explanation of mortality patterns).

As expected given lifespan extension, both H-span and G-span were mainly increased across the 435 treatments (Fig. 2D–E). Notably, the magnitude of G-span increase was greater, attributable to a greater increase in longer-lived population members. Consistent with this was an overall increase in relative G-span (Fig. 2F). These results align with our earlier analysis of 46 treatments from this dataset (4), in which 1. lifespan extension arose primarily from G-span expansion, expanding the proportion of life spent in decrepitude, and 2. G-span expansion was more inter-individually variable than H-span expansion, such that they better explained reductions in, respectively, *β* and *α*. These relationships between overall health, morbidity and lifespan could potentially represent basic biodemographic principles, that are to some degree independent of specific treatments and aging mechanisms.

### Lifespan rectangularization and triangularization reflect distinct changes in health and morbidity

We then asked if these changes in H-span and G-span depend on the mode of lifespan extension: from rectangularizing, triangularizing and non-S-M treatments. First, we examined effects of these three treatment types on lifespan (Fig. 3A); as expected, rectangularizing treatments increased lifespan more in short-lived population members (steepening the survival curve), triangularizing treatments in long-lived population members (flattening the survival curve), and non-S-M treatments produced a mix of these lifespan changes. Interestingly, lifespan shortening by some treatments at the highest and lowest survival proportions (Fig. 2C) is here revealed to reflect lifespan shortening in the longest-lived population members in rectangularizing treatments, and shortest-lived population members in triangularizing treatments (Fig. 3A).

**Fig. 3.**
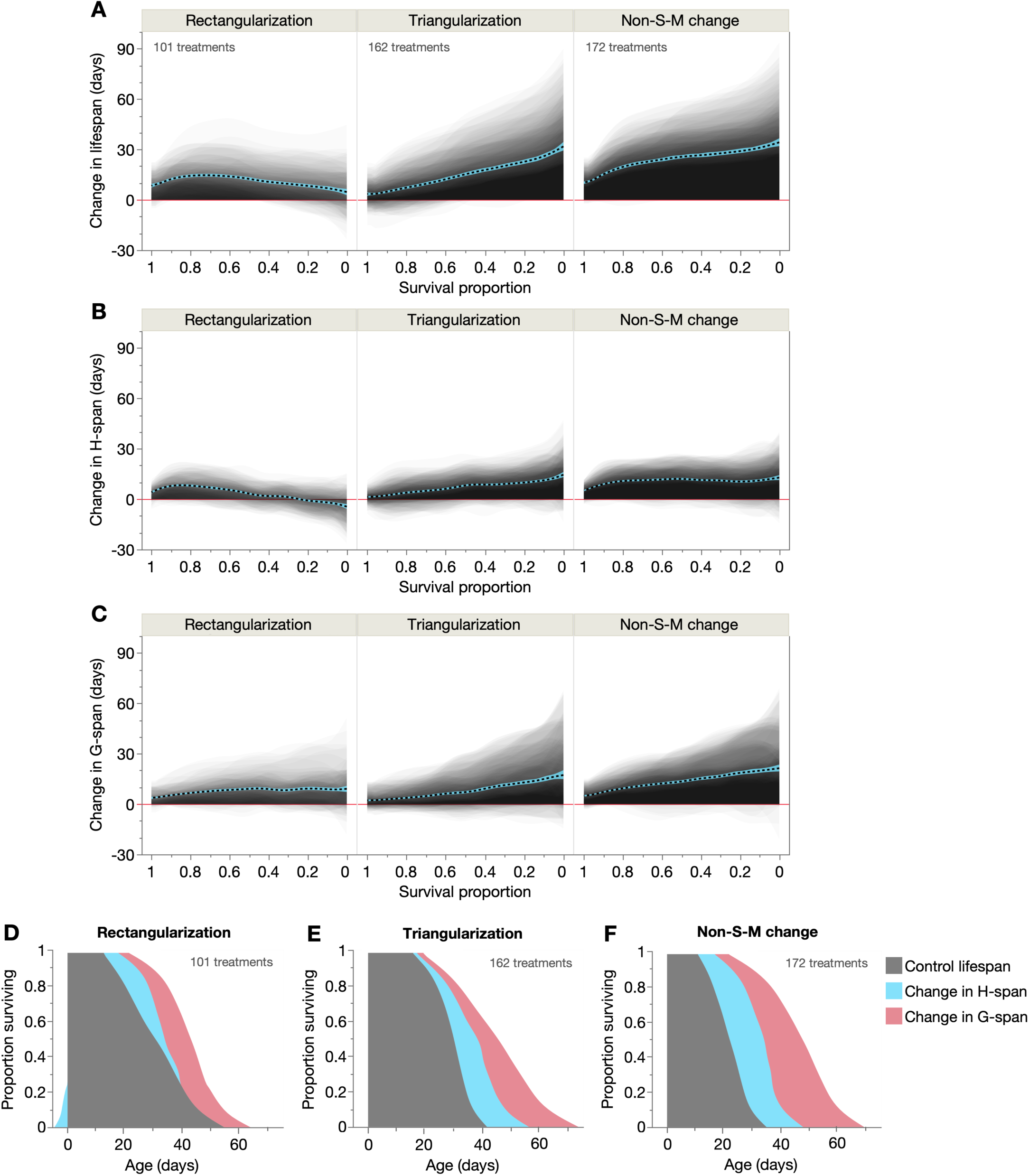
Different demographic life-extension modes have distinct effects on healthspan and gerospan. (**A**–**C**) Shaded and overlaid (JMP transparency=0.02) area plots for all treatments (comparison pairs) of each demographic treatment type (rectangularizing, triangularizing, non-S-M), of the change in (**A**) lifespan, (**B**) H-span, and (**C**) G-span. These shaded changes are plotted over survival proportion (x-axis left: shorter-lived population members, x-axis right: longer-lived population members). Red line: y=0, below which the lifespan-extending treatment shortens that trait. Dashed black line: summary spline smoother (lambda=0.05) of the 435 treatment changes, showing its 95% confidence region in blue. The number of treatments in each demographic type is annotated in **A**. (**D**–**F**) Empirical summary figures of how rectangularizing, triangularizing and non-S-M treatments extend lifespan through additive changes in H-span and G-span (spline smoother of each, stacked on spline smoother of control lifespan), displayed in a shaded survival curve-like format. Shortening of H-span in the longest-lived population members in rectangularizing treatments is represented as a negative change relative to Age=0; this has the effect of further steepening the resultant survival curve.

Mirroring these lifespan changes, H-span was increased by rectangularizing treatments in shorter-lived population members, by triangularizing treatments in longer-lived population members, and by non-S-M treatments relatively evenly across survival proportions (Fig. 3B). Interestingly, H-span was even decreased in the longest-lived population members, explaining their modest lifespan shortening (Fig. 3A). Meanwhile, all three modes of lifespan extension increased G-span primarily in longer-lived population members: moderately in rectangularizing treatments and strongly in triangularizing and non-S-M treatments (Fig. 3C). Notably, these findings were robust to the definition of H-span and G-span, as they were also observed when redefining G-span as only the immotile duration of life (i.e. a stricter G-span definition) (Fig. S4A–B). Additionally, repeated subsampling of biologically-independent treatments (where each cohort is represented only once) yielded the same results, confirming that they are not artifacts of pairwise analysis (Fig. S5A–C). These results reveal distinct combinations of change in H-span and G-span for each mode of lifespan extension, reflecting distinct patterns of change in the biological aging trajectory. Given that each population is isogenic and cultured under identical conditions, this reproducible inter-individual variability of H-span and G-span (e.g. selective extension in short- or long-lived individuals) is particularly striking.

These findings provide a high-level biological view of demographic aging across the 30 cohorts, as individually-resolved distributions of aging-related health and morbidity. These two distributions combine additively to yield the lifespan distribution, thus providing empirical, biological explanations for changes in survival curve shape. These findings are summarised in a shaded survival curve format (Fig. 3D–F). Rectangularizing treatments extend H-span in shorter-lived population members (even truncating it in the longest-lived) and G-span more equally across the population (Fig. 3D). This shows that rectangularization results directly and entirely from changes in H-span (not G-span), because its selective expansion in short-lived population members postpones early mortality, thereby steepening the survival curve. Furthermore, modest H-span shortening in the longest-lived population members truncates the survival curve tail (i.e. maximum lifespan), further steepening and rectangularizing the curve.

In contrast, triangularizing treatments extend both H-span and G-span more in longer-lived population members (Fig. 3E), such that both are responsible for the survival curve tail lengthening that occurs in triangularization. Finally, non-S-M treatments extend H-span equally in all population members and G-span in longer-lived members (Fig. 3F). Here, H-span expansion effectively shifts the survival curve rightward while G-span expansion extends its tail, corroborating our recent findings (4). Notably, these H-span and G-span changes resulting from the non-S-M treatments are intermediate between those of the rectangularizing and triangularizing treatments, consistent with lifespan extension in both short- and long-lived population members.

Subsetting the 435 treatments into those in which only one condition (out of temperature, antibiotic status and genotype) varies at a time yields 30 temperature-only, 15 antibiotic-only, and 60 genotype-only treatments. This showed that temperature-only (i.e. reduced temperature) and antibiotic-only (i.e. carbenicillin) treatments produced all three demographic modes of life-extension, while the genotype-only treatments (i.e. IIS pathway perturbation) selectively produced triangularizing and non-S-M life-extension (Fig. S6). However, the changes in H-span and G-span underlying these three life-extension modes were largely independent of the intervention type (Fig. S6), matching those observed across all 435 treatments (Fig. 3D–F).

### Morbidity expansion and compression by lifespan rectangularization and triangularization

We wondered how the treatments affect the proportion of life spent in age-related morbidity. Notably, all three modes of lifespan extension increased relative G-span (proportion of life in G-span), strongly in rectangularizing and non-S-M treatments and moderately in triangularizing treatments (Fig. 4A, Figs. S4C, S5D). Thus, lifespan extension across these 30 cohorts predominantly causes morbidity expansion rather than compression, and largely irrespective of the demographic mode of longevity. This is consistent with prior reports of morbidity expansion in long-lived *C. elegans* (4, 25–28), although different morbidity measures in certain strains have also suggested proportional morbidity scaling (28, 29).

**Fig. 4.**
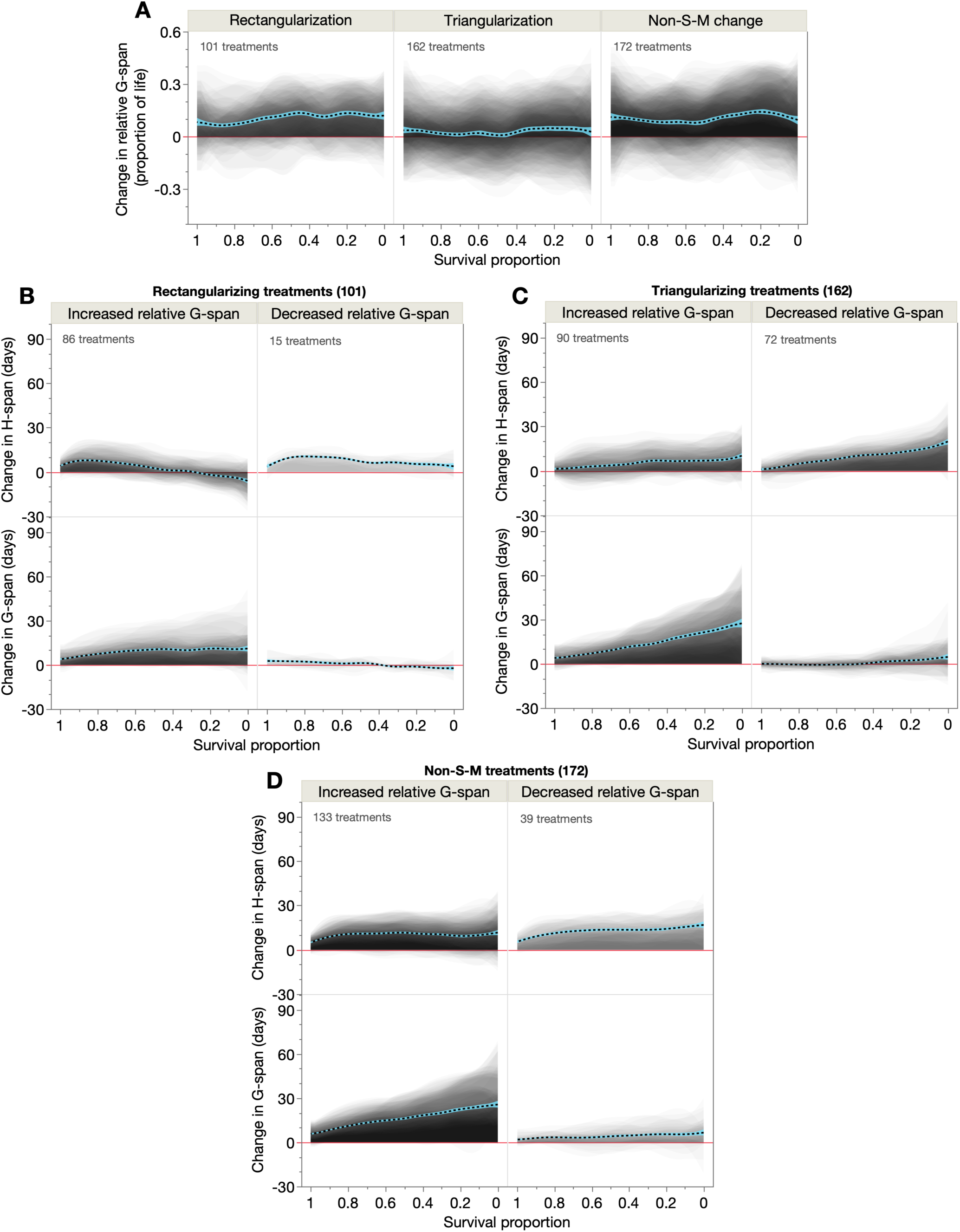
Effects of demographic life-extension modes on morbidity expansion and compression. Shaded and overlaid (JMP transparency=0.02) area plots for all treatments (comparison pairs) of each demographic treatment type (rectangularizing, triangularizing, non-S-M), of the change in (**A**) relative G-span, and (**B**–**D**) H-span and G-span for those subset treatments that increase or decrease relative G-span. These shaded changes are plotted over survival proportion (x-axis left: shorter-lived population members, x-axis right: longer-lived population members). Red line: y=0, below which the lifespan-extending treatment shortens that trait. Dashed black line: summary spline smoother (lambda=0.05) of the changes, showing its 95% confidence region in blue. The number of treatments in each category is annotated.

However, some treatments, especially triangularizing treatments, did decrease relative G-span (Fig. 4A). Such morbidity-compressing treatments are of particular interest; to study them further we partitioned treatments that expanded versus compressed morbidity, for each demographic treatment class (Fig. 4B–D). As expected, most treatments increased relative G-span, via similar demographic changes in H-span and G-span to that observed across all treatments (Fig. 3B–C) but with greater magnitudes of G-span increase.

Meanwhile, the number of rectangularizing, triangularizing and non-S-M treatments that compressed morbidity was, respectively, 15/101 (15%), 72/162 (44%) and 39/172 (23%) (Fig. 4B–D). Thus, most morbidity-compressing treatments (87/126: 69%) are S-M treatments (by comparison, 60% of all treatments are S-M treatments), and of these most (72/87: 83%) are triangularizing treatments. Therefore, within this dataset, triangularizing treatments are best able to compress morbidity while extending lifespan.

How does such morbidity compression arise, in terms of changes in H-span and G-span? The morbidity-compressing rectangularizing treatments increased H-span across all survival proportions, with greatest increases in short-lived population members, while G-span remained largely unchanged (Fig. 4B, right). Similarly, the morbidity-compressing triangularizing treatments robustly increased H-span (in longer-lived population members) but not G-span (Fig. 4C, right). Likewise, the morbidity-compressing non-S-M treatments increased H-span markedly more than G-span (Fig. 4D, right). Compared to the morbidity-expanding treatments (left panels), these morbidity-compressing treatments increased H-span more, and strongly suppressed G-span increases. Thus, these treatments extend lifespan almost entirely through a large H-span extension, such that these individuals experience a postponed *and* brief period of morbidity (Fig. S7A–C).

An interesting question is whether morbidity-compressing treatments involve particular experimental conditions. We therefore asked whether specific temperatures, presence or absence of antibiotic, or specific genotypes were either more or less prevalent than expected by chance, in the control and treatment cohorts of each pairwise treatment class (rectangularizing, triangularizing and non-S-M) (Fig. 5, Table S3–5). Amongst the rectangularizing treatments, genotype, but not temperature or antibiotic usage, was associated with morbidity compression (Fig. 5A–C). Specifically, *daf-2(e1368)* was enriched amongst the treatment cohorts, revealing it to be a major cause of morbidity compression in these rectangularizing treatments. This suggests that unlike other stronger *daf-2* alleles (e.g. *daf-2(e1370)*), *daf-2(e1368)* both maintains wild-type-like levels of voluntary locomotion (as known (27, 28)) *and* exhibits prolonged true locomotory capacity (detected here through stimulated locomotion).

**Fig. 5.**
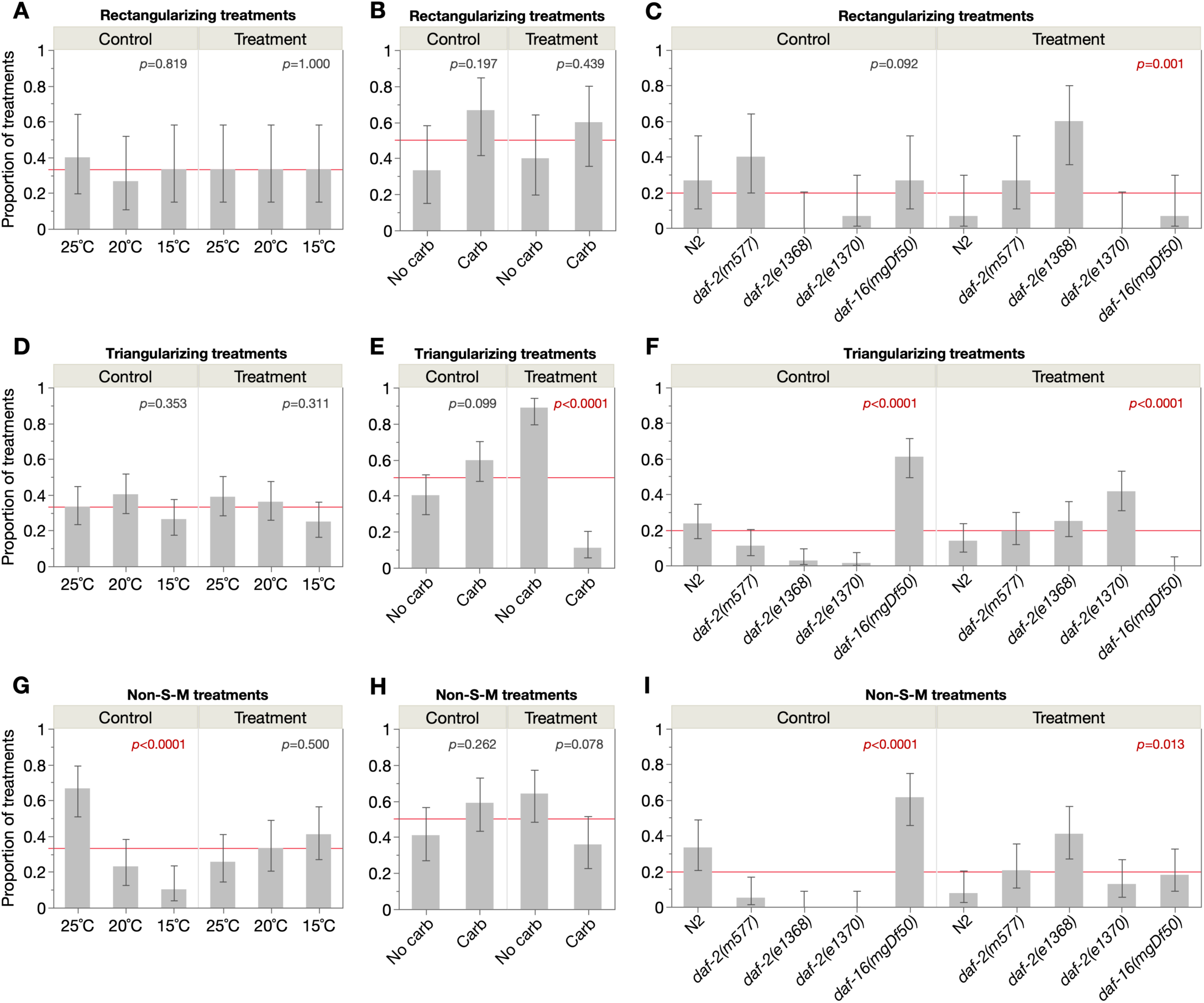
Different demographic life-extension modes compress morbidity through shared and distinct mechanisms. Proportions of the control and treatment cohorts belonging to each experimental condition (3 culture temperatures, ± carbenicillin, 5 IIS-related genotypes), for treatments that decrease relative G-span (i.e. compress morbidity) through (**A**–**C**) rectangularizing, (**D**–**F**) triangularizing or (**G**–**I**) non-S-M demographic changes. Each lifespan-extending treatment comprises a control and treatment cohort, which respectively inform about the mechanistic background and intervention required to compress morbidity. Proportions within each control and treatment panel sum to 1, and proportions above or below the red lines (expected proportions under null hypothesis: 0.33 for 3 temperatures, 0.5 for ± carbenicillin, 0.2 for 5 genotypes) indicate enrichment or depletion of that condition. 95% confidence intervals are shown and Pearson chi-square goodness-of-fit tests were run for each panel (*p* values annotated), with sample sizes of, rectangularization: 15 treatments, triangularization: 72 treatments, and non-S-M change: 39 treatments.

Amongst the triangularizing treatments, both antibiotic usage and genotype were associated with morbidity compression (Fig. 5D–F). This was associated with the *absence* of antibiotic, likely reflecting the morbidity-unmasking effects of preventing life-limiting infection (4, 27) (Fig. 5E). Meanwhile, being enriched amongst control cohorts (Fig. 5F), *daf-16(mgDf50)* was the most responsive genotype to triangularizing morbidity compression (resulting primarily from *daf-2(e1370)* mutation), consistent with the view that mutation of *daf-16* accelerates aging (30, 31) (producing particularly short H-spans). Finally, amongst the non-S-M treatments, cohorts inhabiting higher temperatures (25°C) and again, *daf-16(mgDf50)* cohorts, were particularly responsive to morbidity compression (Fig. 5G–I).

Therefore, these rectangularizing, triangularizing and non-S-M treatments postpone morbidity through common and distinct mechanisms associated with, respectively, IIS, infection and IIS, and temperature and IIS. We also assessed the number of simultaneous condition changes (e.g. 1=temperature change only, 2=temperature and antibiotic usage changes, 3=temperature, antibiotic usage and genotype changes) required for these treatments to compress morbidity. Interestingly, in rectangularizing and triangularizing treatments, 2 condition changes were most common (Fig. S7D), suggesting that interactions between these conditions are required to postpone morbidity, rather than any one alone.

### Rectangularization and triangularization alter the inter-individual variability of aging

Our longitudinal dataset is informative of not only the biological aging trajectory but its inter-individual variability. An important goal of aging research is to explain lifespan variation, both between and within populations; the latter is particularly intriguing in isogenic and environmentally-controlled populations, as in this dataset. In *C. elegans*, lifespan variation is particularly attributable to inter-individual variation in the duration of late-life morbidity (4, 22, 28, 32), and correlates with early life molecular and cellular traits (33–37). Therefore, we wondered whether the decreased and increased lifespan variation, respectively, in survival curve rectangularization and triangularization is determined by similar variability changes in the preceding biological aging process (e.g. in H-span and G-span, amongst other traits).

As expected, lifespan standard deviation was decreased in rectangularizing treatments and increased in triangularizing treatments (Fig. 6A, left). This reflects the compression of mortality and extension of maximum lifespan, respectively, that occur in these demographic changes (Fig. 1C, E, F, H). The non-S-M treatments also increased lifespan standard deviation, but less so than triangularizing treatments, consistent with increases in both maximum and early lifespan.

**Fig. 6.**
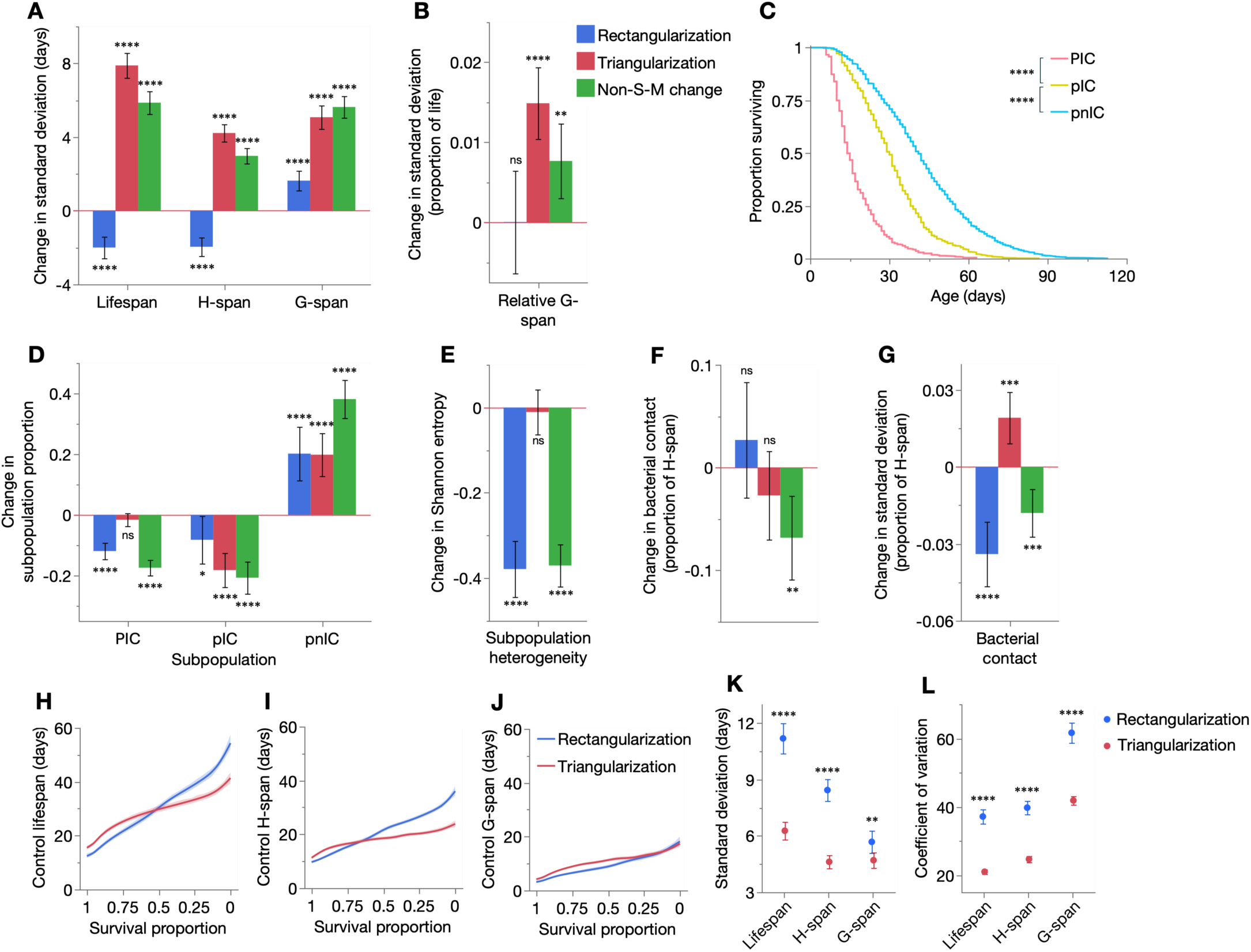
Rectangularizing and triangularizing treatments, respectively, homogenize and heterogenize the aging process. (**A**–**B**) Mean changes in standard deviation of (**A**) lifespan, H-span, G-span and (**B**) relative G-span, for rectangularizing, triangularizing and non-S-M treatments. (**C**) Kaplan-Meier survival curves of PIC, pIC and pnIC individuals from all 30 cohorts (*n*=443, 991, 2144, respectively); censored individuals were excluded. Lifespans were compared statistically using the log-rank test. P: infected swollen pharynx; p: uninfected pharynx (20). Therefore, PIC: <u>P</u> with intestinal <u>c</u>olonisation, pIC: <u>p</u> with intestinal <u>c</u>olonisation, pnIC: <u>p</u> with <u>n</u>o intestinal <u>c</u>olonisation (4). (**D**) Mean changes in subpopulation proportion by rectangularizing, triangularizing and non-S-M treatments, and (**E**) mean changes in population heterogeneity resulting from these subpopulation proportion changes, as measured by Shannon entropy. (**F**–**G**) Mean changes in the (**F**) mean and (**G**) standard deviation of bacterial contact (proportion of H-span in contact with *E. coli* lawn) by rectangularizing, triangularizing and non-S-M treatments. (**H**–**J**) Spline smoother of mean (**H**) lifespan, (**I**) H-span and (**J**) G-span of control cohorts that responded to life-extending treatments with rectangularization versus triangularization, plotted over survival proportion (x-axis left: shorter-lived population members, x-axis right: longer-lived population members). 95% confidence regions are shaded. (**K**–**L**) Mean (**K**) standard deviation and (**L**) coefficient of variation of lifespan, H-span and G-span of control cohorts that responded with rectangularization versus triangularization (*n*=101, 162). Means were compared statistically using two-tailed Student’s t-tests; 95% confidence intervals are shown. Changes in means in **A**–**B** and **D**–**G** were compared statistically using two-tailed one sample t-tests (*H*_0_: mean change=0), showing 95% confidence intervals; Benjamini-Hochberg correction of *p* values did not change which treatments reached the 0.05 significance threshold (data not shown). In all panels: ns *p* > 0.05, * *p* ≤ 0.05, ** *p* ≤ 0.01, *** *p* ≤ 0.001, **** *p* ≤ 0.0001.

Notably, inter-individual variation in H-span and G-span changed in the same directions as lifespan variation, except for an increase in G-span variation in rectangularizing treatments (Fig. 6A, Fig. S8A–B). In line with these changes, variation in relative G-span was unaltered in rectangularizing treatments and increased in triangularizing and non-S-M treatments (Fig. 6B, Fig. S8D–E). Normalisation of these changes to their means (coefficient of variation) revealed that respectively, rectangularizing and triangularizing treatments disproportionately decreased and increased inter-individual variation in H-span, G-span and relative G-span (Fig. S8C, F). These findings suggest that rectangularizing and triangularizing treatments respectively homogenize and heterogenize the aging process, resulting in respectively reduced and increased inequalities in the durations of health and morbidity.

These changes in H-span and G-span may reflect changes in biological aging rate across the life course, which would stretch or compress their durations. However, individual *C. elegans* vary not only in the speed of aging but also its trajectory. Distinct (yet isogenic) subpopulations have been characterised, which acquire aging-related, organ-specific colonizations by the dietary *E. coli*, affecting both the pharynx and intestine (“PIC” individuals), the intestine only (“pIC” individuals), or neither organ (“pnIC” individuals) (4, 20, 38) (Fig. 6C caption). These subpopulations were scored in the present dataset, by necropsies on 91% of all uncensored individuals (3,578/3,911) across the 30 cohorts. Notably, the three subpopulations had different longevities, with longer lifespans when fewer organs were bacterially colonized (Fig. 6C).

Notably, rectangularizing treatments primarily reduced the prevalence of the short-lived PIC individuals (and intermediate-lived pIC individuals), replacing them with individuals from the longest-lived pnIC subpopulation (Fig. 6D). Thus, lifespan rectangularization here results from the redirection of individuals from shorter- to longer-lived aging trajectories, which postpones and compresses mortality into the survival curve tail. Conversely, triangularizing treatments converted only pIC individuals into pnIC individuals (Fig. 6D), thus triangularizing the survival curve by maintaining early mortality while pushing maximum lifespan. Finally, the non-S-M treatments converted both PIC and pIC individuals to pnIC, consistent with their postponement of both early and late mortality. Therefore, these three demographic modes of lifespan extension involve distinct individual-specific changes in both the rate and trajectory of aging.

Notably, rectangularizing treatments decreased two measures of subpopulation heterogeneity (Shannon entropy and Simpson’s Diversity Index, applied to PIC, pIC and pnIC proportions) (Fig. 6E, Fig. S8J), supporting their action as homogenizing treatments. However, triangularizing treatments did not change these measures of subpopulation heterogeneity, suggesting they may increase inter-individual variation in the rate (H-span and G-span) but not trajectory of aging.

We also asked how early in life such variability changes emerge. Given aging-related colonization by dietary *E. coli*, and that intestinal *E. coli* load in early adulthood already predicts nematode lifespan (33), we longitudinally scored the proportion of H-span spent in contact with the bacterial lawn, for all individuals of the 30 cohorts in at least 3 trials. Mean bacterial contact of cohorts was largely unchanged by rectangularizing and triangularizing treatments, and decreased by non-S-M treatments (Fig. 6F). Interestingly, however, inter-individual variation in bacterial contact was mostly decreased by rectangularizing treatments and increased by triangularizing treatments (Fig. 6G, Fig. S8G–I), consistent with the dominant directions of variation change in H-span, G-span, relative G-span, disease subpopulation heterogeneity and lifespan. Taken together, these findings demonstrate that inverse Gompertz parameter changes reflect fluctuations in the inter-individual variability of aging: homogenization in rectangularization and heterogenization in triangularization. Consistent with this, the non-S-M treatments, which produce intermediate changes in lifespan variation (between rectangularization and triangularization), exhibited greater variation in the type of variability change seen (i.e. homogenization or heterogenization).

Finally, we wondered what determines whether a given life-extending intervention causes rectangularization or triangularization. Intriguingly, lifespan, H-span and G-span were consistently more inter-individually variable in the control cohorts that underwent rectangularization than triangularization (Fig. 6H–L). This argues that demographic responses to lifespan-extending interventions depend on the existing level of variation: heterogeneous populations are more likely to rectangularize, and homogeneous populations to triangularize. Through which specific biological determinants this demographic decision is made remains to be uncovered.

### Conservation of rectangularization and triangularization determinants in *Drosophila* and mice

Finally, we wondered whether we could explain the biological basis of rectangularization and triangularization in other species. Species differences in the aging process and in health/morbidity definitions may argue against the conservation of nematode H-span/G-span dynamics across species. However, these dynamics could potentially be upstream of specific mechanisms and thereby follow similar biodemographic principles.

Availability of lifelong longitudinal health data of other animal models is more limited than for *C. elegans*. We analysed two *Drosophila* and three mouse datasets, which despite having smaller cohort sizes and often less frequent longitudinal health measurements, importantly contained health and lifespan data from the same individuals, across multiple cohorts. We therefore similarly treated each lifespan-extending pairwise comparison between cohorts as a lifespan-extending treatment, categorized them as rectangularizing or triangularizing treatments, and compared their effects on lifespan, H-span, G-span and relative G-span.

The *D. melanogaster* datasets measured age-related locomotory decline in 14 cohorts undergoing different dietary regimes (protein:carbohydrate ratio, and curcumin and fruit extract supplementation; both sexes) (39), and reproductive aging (age-decline in fecundity) in females of 9 laboratory-cultured and wild-caught lines meta-analytically compiled from several primary studies (40) (Table S6, 7). In both datasets, rectangularizing treatments primarily increased H-span, in short- to intermediate-lived population members (Fig. 7A–B, D–E). G-span was decreased in the longest-lived individuals, but only modestly. Thus, as in our nematode cohorts, it is primarily H-span expansion that rectangularizes the survival curve. Also similar to the nematode data, the triangularizing treatments generally increased H-span and G-span in longer-lived population members (except H-span in the reproductive aging data) (Fig. 7A, C, D, F). Furthermore, in both datasets, relative G-span increased marginally in triangularizing treatments; however, the rectangularizing treatments trended towards morbidity compression (Fig. 7A, D, bottom). The biological determinants of these survival curve transformations in *Drosophila* therefore overlap with those in *C. elegans*.

**Fig. 7.**
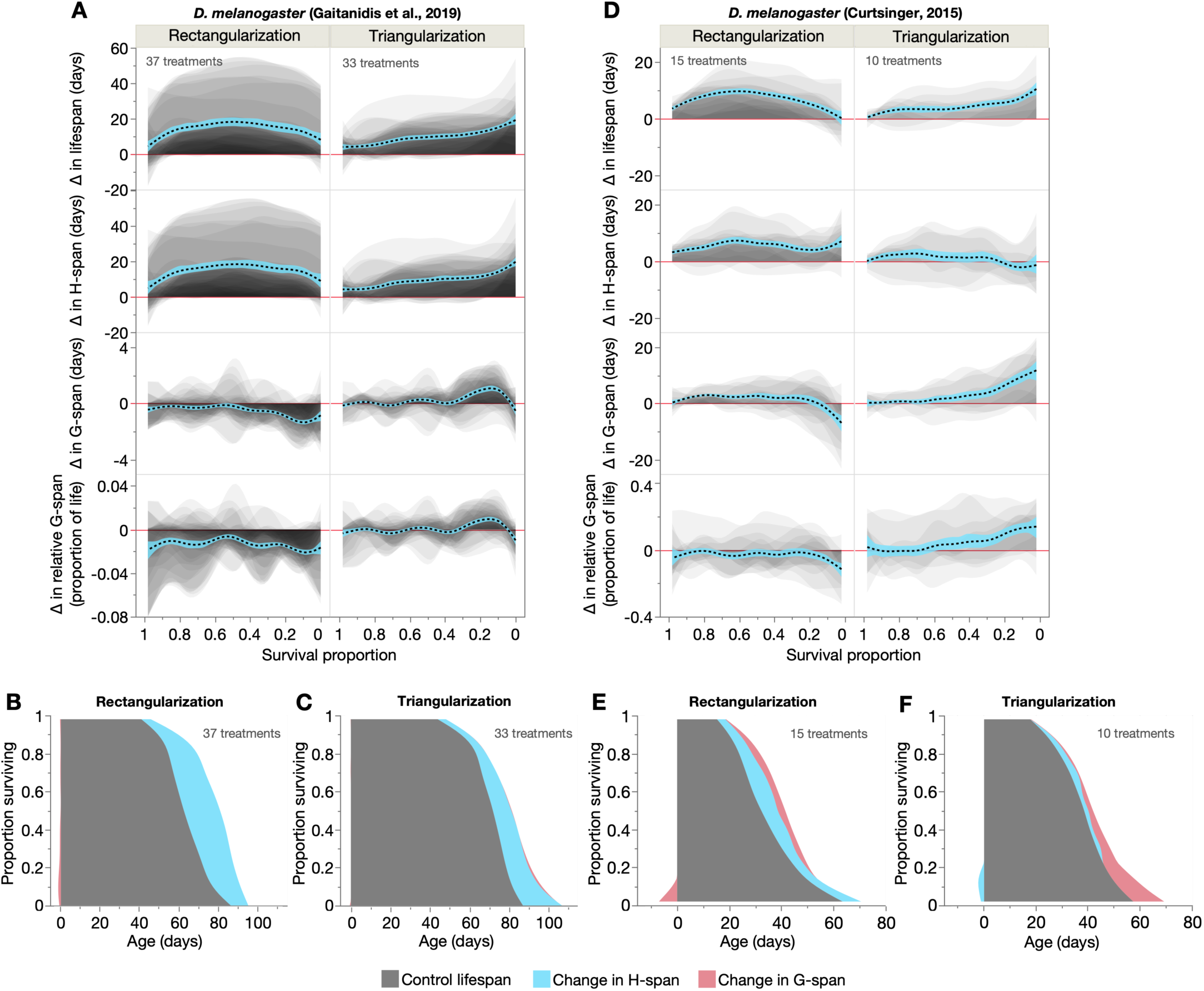
Health and morbidity profiles of rectangularization and triangularization in *Drosophila melanogaster*. (**A**, **D**) Shaded and overlaid (JMP transparency=0.05) area plots for lifespan-extending rectangularizing and triangularizing treatments, of changes in lifespan, H-span, G-span and relative G-span. These shaded changes are plotted over survival proportion (x-axis left: shorter-lived population members, x-axis right: longer-lived population members). Red line: y=0, below which the lifespan-extending treatment shortens that trait. Dashed black line: summary spline smoother (lambda=0.05) of the treatment changes, showing its 95% confidence region in blue. The number of rectangularizing and triangularizing treatments is annotated. (**B**–**C**, **E**–**F**) Empirical summary figures of how rectangularizing and triangularizing treatments extend lifespan through additive changes in H-span and G-span (spline smoother of each, stacked on spline smoother of control lifespan), displayed in a shaded survival curve-like format. Shortening of H-span and G-span is represented as a negative change relative to Age=0.

We next examined the three mouse cohorts, which measured longitudinal age changes in a 32-item fragility index in response to dietary restriction and intermittent fasting (41) (7 cohorts), and 31-item frailty indexes given alpha-ketoglutarate treatment (42) (4 cohorts) and mutation of RNA polymerase III (*Polr3b^+/-^*) (43) (4 cohorts; previously unpublished frailty data) (Table S8–10). The first dataset yielded primarily triangularizing treatments (Fig. 8A), consistent with reports of *β* reduction by dietary restriction in rodents (44, 45), while the latter two datasets yielded rectangularizing treatments (Fig. 8D, F). In all three datasets, rectangularizing treatments primarily increased H-span, which occurred mainly in short-lived population members (Fig. 8A–B, D–G), thus rectangularizing the survival curve as in the nematode and *Drosophila* datasets. However, in the dietary restriction dataset, H-span also increased in longer-lived population members and G-span increased mildly in the shortest-lived, suggesting that here rectangularization arises from combined changes in H-span and G-span (Fig. 8A–B). Survival curve triangularization was driven alone by G-span increases in longer-lived population members (Fig. 8A, C). Across these datasets, relative G-span was either unchanged or increased, consistent with the greater frequency of morbidity expansion in our nematode treatments.

**Fig. 8.**
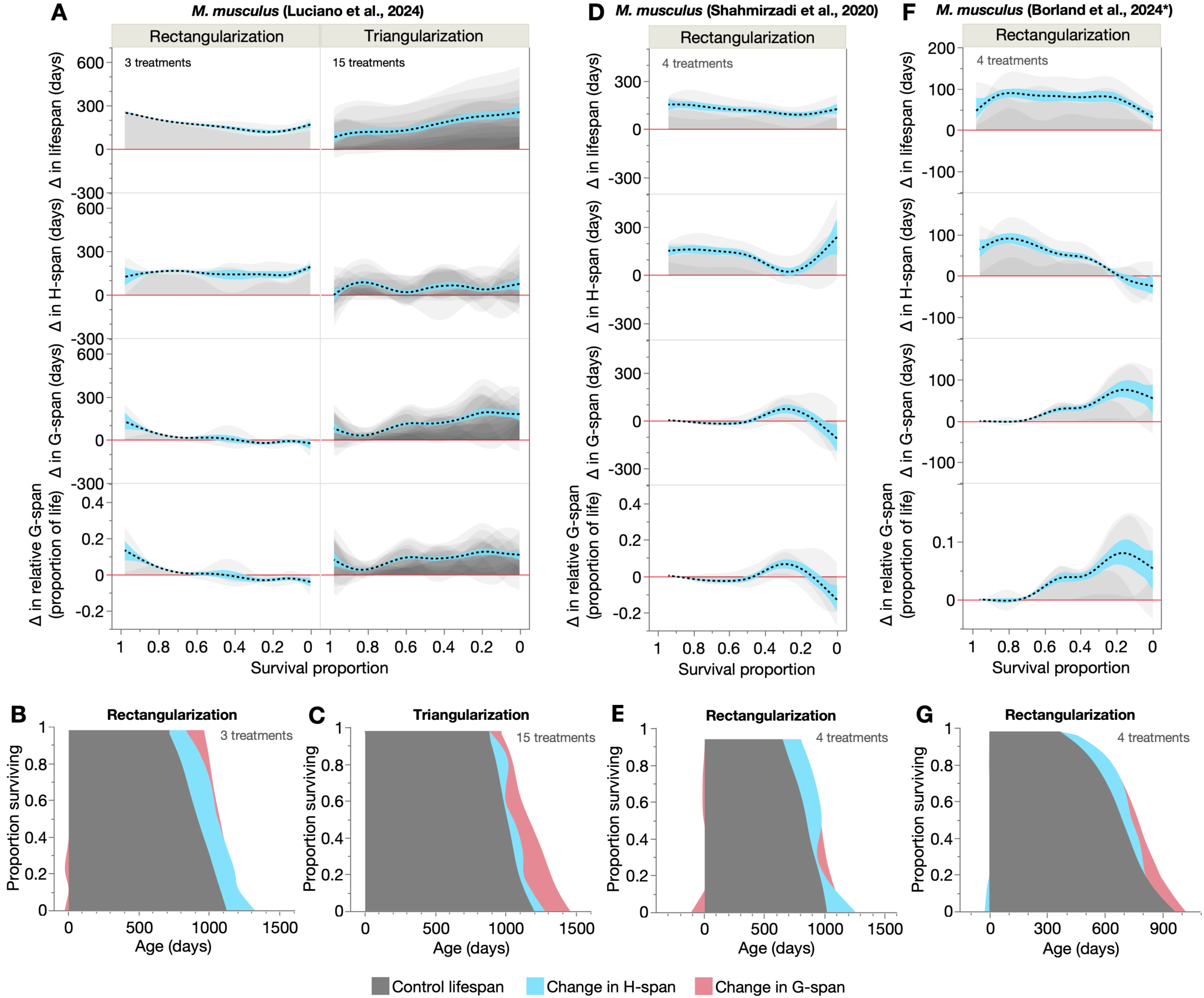
Health and morbidity profiles of rectangularization and triangularization in mice. (**A**, **D**, **F**) Shaded and overlaid (JMP transparency=0.05) area plots for lifespan-extending rectangularizing and triangularizing treatments, of changes in lifespan, H-span, G-span and relative G-span. These shaded changes are plotted over survival proportion (x-axis left: shorter-lived population members, x-axis right: longer-lived population members). Red line: y=0, below which the lifespan-extending treatment shortens that trait. Dashed black line: summary spline smoother (lambda=0.05) of the treatment changes, showing its 95% confidence region in blue. The number of rectangularizing and triangularizing treatments is annotated. (**B**–**C**, **E**, **G**) Summary figures of how rectangularizing and triangularizing treatments extend lifespan through additive changes in H-span and G-span, displayed in a shaded survival curve-like format. Shortening of H-span and G-span is represented as a negative change relative to Age=0. **\***Previously unpublished frailty data (**F**–**G**), from the mouse cohorts in Borland et al. (2024) (43).

Taken together, these nematode, fruit fly and mouse cohorts exhibit both common and distinct biological bases of rectangularization and triangularization. Despite species differences in the aging process and H-span/G-span definitions, rectangularization of survival curves was driven by H-span expansion in shorter-lived population members, and triangularization by G-span expansion in longer-lived population members. These similarities may reflect conserved demographic dynamics of how durations of health and morbidity vary between individuals, and respond to lifespan-extending interventions. Indeed, strikingly, as in our nematode dataset (Fig. 6H–L), lifespan variation was consistently higher in the control cohorts that underwent rectangularization than triangularization, in the two *Drosophila* and one mouse datasets that contained both modes of life-extension (Fig. S9). This suggests another species-independent principle of demographic lifespan extension, wherein heterogeneous populations undergo rectangularization and homogenous populations triangularization.

## Discussion

Here, we used *C. elegans*, *D. melanogaster* and mice to empirically determine the biological basis of lifespan-extending, inverse changes between parameters of the Gompertz mortality model. Through the simultaneous longitudinal study of individual and population aging across conditions and species, we could resolve the biological states underlying the rectangularization and triangularization of survival curves (which occur when Gompertz parameters change inversely). Rectangularization reflected healthspan changes in shorter-lived population members, whereas triangularization reflected a combination of healthspan and gerospan changes in longer-lived population members (Fig. 9). Importantly, rectangularization and triangularization respectively homogenized and heterogenized the biological aging process between population members, depending on the degree of existing heterogeneity (Fig. 9).

**Fig. 9.**
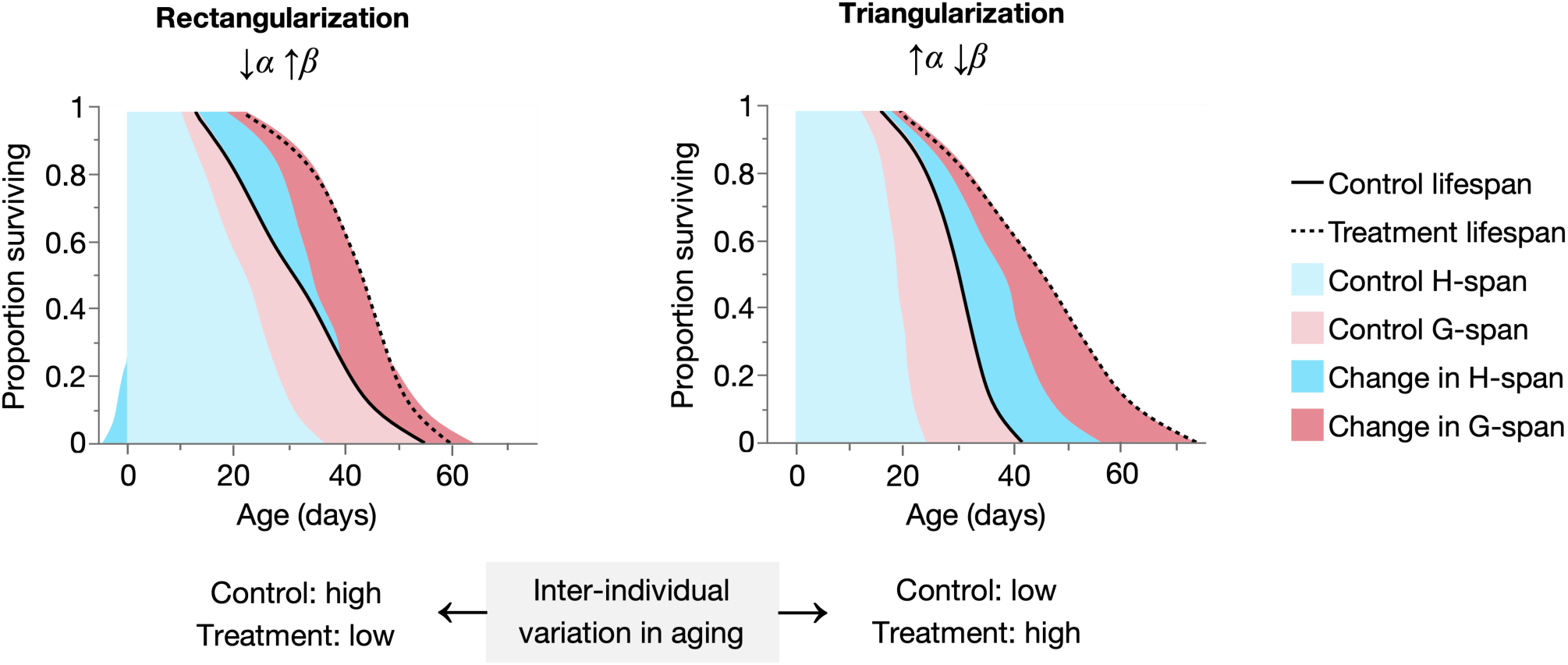
Summary schematic: rectangularization and triangularization decrease and increase variation in aging, respectively. Empirical summary figures depicting how rectangularization and triangularization extend lifespan through additive changes in H-span and G-span, in a shaded survival curve-like format. Rectangularization results from increases in H-span in shorter-lived population members, accompanied by relatively equal G-span increases across the population, whereas triangularization results from increases in both H-span and G-span in longer-lived population members. These changes (and others; see Fig. 6, Fig. S8) reflect decreases and increases, respectively, in the inter-individual (within-population) variability of the aging process by interventions that rectangularize and triangularize the survival curve. Notably, such interventions cause rectangularization when existing variation (in control cohorts) is high (thereby reducing it), and triangularization when the existing variation is low (thereby increasing it).

Given that rectangularization and triangularization are the survival curve transformations that occur in Strehler-Mildvan correlations, our empirical explanation of these transformations may potentially also explain the S-M correlation. S-M correlations (measured across multiple populations) typically reflect a continuum of rectangularization or triangularization across several populations (e.g. Fig. 1B), suggesting that they all vary primarily (but to different degrees) in some same, complex biological state or process. Across our 30 nematode populations however, an overall S-M correlation was not observed, likely reflecting the diversity of interventions applied and combined, which affect different mechanisms in the aging process. Yet the many instances of rectangularization and triangularization within this dataset, despite resulting from biologically-diverse interventions, show the same biological underpinnings – supporting their likely relevance to datasets displaying overall S-M correlations. This could be tested by selecting any intervention that rectangularizes or triangularizes survival, generating several populations with differing ‘dosages’ of the intervention, and if they yield an S-M correlation between them, to assay the individual biologies of all populations’ members.

Our longitudinal health data on rectangularization and triangularization may be instructive for the validation and development of theoretical aging models that predict the S-M correlation (5, 12, 46). The data also provide an account of the biology underlying non-S-M-type changes, where lifespan extension occurs through reductions in both Gompertz parameters.

Our findings by extension favour inter-individual heterogeneity explanations of the S-M correlation over those assuming intra-individual trade-offs between health and aging (13–15). Also supporting heterogeneity explanations, reduction of biological noise in a mathematical model of aging corresponded to lifespan rectangularization, as supported by human and canine data (47, 48). Additionally, decreasing a noise parameter in another mathematical aging model rectangularized survival, although this was predicted to compress relative gerospan, contrasting against our empirical observation of morbidity expansion in rectangularization (49). However, in agreement, increasing the death threshold parameter of this model also rectangularized survival and did predict morbidity expansion (49).

Meanwhile, trade-off explanations of S-M correlations often arise from theoretical interpretations of model parameters, such as the conventional equation of *α* and *β* with aging-independent processes and biological aging rate, respectively (50, 51). However, our work argues against these interpretations (in fact, inverting them (4)); indeed, that healthspan extensions cause lifespan rectangularization is a further argument against traditional interpretations of increased *β* as accelerated aging. We also show that within isogenic populations, biological aging rates and trajectories vary greatly. Mortality patterns may therefore be more appropriately studied as inter-individual *distributions* of biological aging processes. Understanding such variability distributions, which though seemingly stochastic in origin (52), are highly reproducible and likely to be foundational for understanding and intervening in aging.

Why are rectangularization and triangularization (and the S-M correlation) so common across conditions and species? Our findings suggest that inter-individual variation, present in all populations, controls fundamental biodemographic dynamics of mortality. In all species examined (nematodes, flies and mice), the amount of cohort heterogeneity predicted whether lifespan-extending interventions would rectangularize or triangularize survival. This may reflect the greater likelihood of homogeneous populations to lose (rather than further increase) homogeneity, and vice versa for heterogeneous populations, somewhat akin to the principle of regression towards the mean (53). Additionally, both rectangularization and triangularization involve changes that affect individual longevity to different extents, consistent with known individual-specificities of many lifespan- and health-modulating interventions (54, 55).

We observed both similarities and differences in the healthspan and gerospan dynamics underlying rectangularization and triangularization between nematodes, flies and mice. In common was healthspan expansion in shorter-lived population members in rectangularization, and gerospan expansion in longer-lived population members in triangularization, although these patterns varied in magnitude. Such cross-species comparisons are limited by data availability and resolution (e.g. population size and health measurement frequency), and complicated by differing health/morbidity definitions, though larger longitudinal studies capturing multiple aging phenotypes are underway (56, 57). In particular, evaluating biodemographic determinants using different definitions of healthspan and gerospan would be important considering the multifactorial complexity of aging.

Are our findings relevant to human populations? Human lifespan has rectangularized over the last two centuries due largely to reductions in early-life mortality by communicable diseases (58), concentrating deaths into later ages (59, 60). This may extend lifespan mainly through healthspan increases in shorter-lived population members, by preventing premature death in these otherwise healthy individuals. Notably we observed this, as shorter-lived, bacterially-infected subpopulations were converted into longer-lived, uninfected subpopulations, in our rectangularizing nematode treatments.

Importantly, the homogenization of human populations (towards aging-related deaths) has slowed further reductions in early mortality (61), while maximum lifespan has gradually increased in more recent decades, reflecting medical advances against aging-related diseases (61–63). In fact, decelerations of mortality rate increase at advanced ages have been observed (64, 65) (although this remains debated (66, 67)), as well as increased inter-individual variation in human health (68). Intriguingly, these changes are reminiscent of lifespan triangularization, suggesting that human survival may be experiencing a demographic transition from rectangularization to triangularization. Supporting this possibility, we observed that lifespan extension of homogeneous (rectangularized) nematode, fly and mouse populations occurs through subsequent triangularization. Our triangularizing nematode treatments increased both healthspan and gerospan in the survival curve tail, and produced the smallest increases in relative morbidity, compared to the other modes of lifespan extension. Whether similar health and morbidity dynamics could underly human triangularization remains to be seen.

## Materials and Methods

### *C. elegans* culture and strains

*C. elegans* were maintained at 20°C using standard protocols (69), on Nematode Growth Medium (NGM, containing Bacto Peptone) plates seeded 2 days before use with a bacterial food source (*Escherichia coli* OP50). Floxuridine (5-fluoro-2-deoxyuridine), sometimes used to block progeny production, was not used in this study. Nematode strains used were: N2 (wild type, hermaphrodite stock (70)), GA1959 *daf-2(m577) III*, GA1960 *daf-2(e1368) III*, GA1928 *daf-2(e1370) III*, and GA1952 *daf-16(mgDf50) I*. All strains were raised from egg at 20°C on live *E. coli*, and transferred at L4 stage to the appropriate experimental conditions (15°C, 20°C or 25°C; with or without carbenicillin). Carbenicillin solution was added topically to plates one day before adding animals (further details below).

### Aging cohorts and lifespan scoring

Nematodes were cultured throughout life in 60 mm Petri dishes (containing 10 mL of NGM) seeded with approximately 80 μL of *E. coli*, and where relevant, treated with 80 μL of 500 mM carbenicillin (Fisher Scientific Ltd, catalogue no. 12737149). In each trial, at L4 stage (time 0 in all analyses), 25 animals were placed on a plate, with two plates per condition. Animals were transferred every two days during the reproductive period. Following the end of egg laying, animals were transferred to individual wells of 24-well tissue culture plates, containing 2 mL of NGM and seeded with 3.5 μL of *E. coli* OP50, and where relevant, treated with 16 μL of 500 mM carbenicillin. Animals were subsequently transferred to fresh 24-well plates monthly, before media desiccation (plates were sealed with parafilm to delay desiccation, and to prevent bacterial/fungal contamination). Scoring of survival was performed every 2–3 days alongside scoring of locomotory class, and necropsy at death (described below). Animals showing no movement were gently touched with a platinum wire (worm pick) on the head and/or tail; those that showed no movement at all in response were scored as dead. Animals that died due to desiccation on the Petri dish wall, internal hatching of larvae, or rupture of internal tissues through the vulva, or that became contaminated by non-*E. coli* bacteria or fungi, or could not be found, were censored.

### Quantification of locomotory decline with age

Locomotory health class (belonging to H-span or G-span) was scored by classifying individuals into one of three classes, adapted from earlier systems (23, 24): A – sinusoidal locomotion; B – non-sinusoidal locomotion; C – no locomotion. To accurately determine locomotory class, animals were gently touched on the tail with a platinum wire worm pick for up to 20 seconds to induce an escape response that reveals movement capacity (rather than behavioral preference (29)), and additionally on the head as a final check. The duration spent in A class was defined as H-span, and the summed duration spent in B and C classes as G-span. Here, B and C classes were summed to improve data tractability, and to provide a definition of G-span that captures both early and late-stage functional declines. However, an alternative definition of the summed duration in A and B classes as H-span, and duration in only C class as G-span, was also tested (Fig. S4), which yielded the same conclusions.

### Necropsy analysis

Necropsy to define patterns of *E. coli*-associated pathology was performed by examining fresh corpses under a Leica MZ8 stereomicroscope (50x magnification). Scoring of swollen, bacterially-infected pharynxes (P), and uninfected, atrophied pharynxes (p) was performed as previously described (20). Intestinal colonization (IC) with *E. coli* was scored where severe bacterial accumulation was observed in the anterior and/or posterior intestine. Such colonization presented as extreme lumenal distension by proliferating bacteria and/or colonization of the intestine beyond the lumenal barrier, with concomitant intestinal tissue degeneration. In pharyngeal and intestinal tissues, sites of bacterial colonization exhibit a yellow-brown colour (like that of the *E. coli* lawn and colocalizing with RFP-labelled *E. coli*), translucent and uniform texture (loss of healthy tissue structures that otherwise appear dark, granular and opaque), and swollen/distended morphology (extensive proliferation of live *E. coli*) (4).

### Bacterial contact scoring

Bacterial contact was scored every 1–2 days. Animals were scored as in contact with bacteria if they were observed inside the lawn at first sighting; each animal was checked once during each scoring session. Importantly, plates were handled carefully to avoid sudden movements or knocks that could startle animals and affect their location within their wells. Notably, animals were often observed to be stationary at the edge of the lawn, resting with their heads outside of the lawn; these animals were scored as not in contact with the bacteria. Accordingly, animals resting with their heads inside the lawn but tail outside were scored as in contact with the bacteria. Where nematode tracks led to growth of *E. coli* outside of the central lawn, nematode interaction with these colonies were treated the same as for the central lawn. If the media surface approached complete coverage by *E. coli*, animals were transferred to fresh wells to allow detection of bacterial avoidance.

For analysis, bacterial contact was quantified as the proportion of H-span (duration in A class) spent in contact with the bacteria, to avoid confounding effects of locomotory decline on the locomotion-dependent bacterial contact phenotype. Because scoring commenced in most cases after the reproductive period (when animals were moved to 24-well plates), H-span was here modified by subtracting the number of days during which bacterial contact was not scored.

### Statistics, software and data handling

Lifespan, locomotory and necropsy data for N2, *daf-2(m577)*, *daf-2(e1368)* and *daf-2(e1370)* cohorts have been analyzed elsewhere (4), but not data for *daf-16(mgDf50)* cohorts and bacterial contact of all cohorts. Statistical tests and figure construction were performed in JMP Pro (SAS Institute, Inc.), except for Gompertz parameter estimation and assessment of statistical differences between them, which were performed using WinModest (19). Right censors were included in all Kaplan-Meier and WinModest analyses, and excluded from others. Specific statistical tests and associated methodological details are described in the respective figure/table captions. Notation of statistical significance in all figures is as follows: * *p* ≤ 0.05, ** *p* ≤ 0.01, *** *p* ≤ 0.001, **** *p* ≤ 0.0001. The measure of subpopulation heterogeneity, Shannon entropy, was calculated as –([PIC proportion ⋅ log(PIC proportion)] + [pIC proportion ⋅ log(pIC proportion)] + [pnIC proportion ⋅ log(pnIC proportion)]), and an alternative measure, Simpson’s Diversity Index, was calculated as 1 – [(PIC proportion)^2^ + (pIC proportion)^2^ + (pnIC proportion)^2^].

### Drosophila data

*Drosophila melanogaster* data were obtained from two studies: Gaitanidis et al. (2019) (39) (the processed data is publicly available in and was obtained from Yang et al. (2025) (49)) and Curtsinger (2015) (40). H-span and G-span in our reanalysis corresponds to “health-span” and “ill-span” in Gaitanidis et al. (2019). The Curtsinger (2015) data are a meta-analysis (40) of the data from several earlier studies (71–74). These were extracted from Figure S3 using WebPlotDigitizer (75), producing 100 ‘pseudoindividuals’ that are representative of their survival proportion value (actual cohort sizes are summarized in Curtsinger (2015)), where H-span and G-span respectively correspond to the number of days of life with at least one, or zero eggs laid. Days before day 8 were excluded due to low fecundity arising from reproductive immaturity rather than aging. In all datasets, the direction of changes in lifespan, H-span and G-span was always calculated in the lifespan-increasing direction (i.e. change = longer-lived cohort – shorter-lived cohort).

### Mouse data

Mouse data were obtained from three sources: Luciano et al. (2024) (41), Shahmirzadi et al. (2020) (42), and previously unpublished frailty data from the mouse cohorts from Borland et al. (2024) (43). In these datasets, G-span onset was defined as the first age at which fragility or frailty index reached or exceeded a threshold value: 0.25 (fragility index) for Luciano et al. (2024), and 0.2 (frailty index) for Shahmirzadi et al. (2020) and Borland et al. (2024) cohorts, when a spline smoother (lambda=0.05) was fit through each individual’s fragility/frailty index values, starting from a theoretical 0 value at 100 days (approximate age of maturity, prior to aging). The thresholds were selected to reflect a notable number of simultaneous aging-related deficits (i.e. not too low) and to maximise the number of individuals that reached it within their lifetimes (i.e. not too high). In the data from Luciano et al. (2024), individuals that went missing, failed to recover from anaesthesia, or were discarded were included in survival and Gompertz analysis as right censors. In all datasets, the direction of changes in lifespan, H-span and G-span was always calculated in the lifespan-increasing direction (i.e. change = longer-lived cohort – shorter-lived cohort).

The mouse cohorts from Borland et al. (2024) were maintained as previously described (43). Frailty index measurements were taken for all mice at 12, 18, 21, 23, 25, 27, 29 and 31 months of age, using a 31-point frailty index system (76). Experimental procedures were approved by The University of Glasgow Animal Welfare and Ethical Review Board, under a UK Home Office Project Licence (PDBDC7568) and following the “principles of laboratory animal care” (NIH Publication No. 86–23, revised 1985).

## Acknowledgments

Some nematode strains were provided by the Caenorhabditis Genetics Center, which is funded by the NIH Office of Research Infrastructure Programs (P40 OD010440). We are grateful to Johnathan Labbadia and Peter Fedichev for helpful discussion, Jennifer M.A. Tullet and Nazif Alic for contributing to the generation of knowledge, and to Stephen E. Wilkie, Zhe Wang and the Biological Services Staff (University of Glasgow) for help with mouse data collection and animal husbandry. The mouse line C57BL/6N-Polr3b/Tcp was generated as part of the NorCOMM2 project funded by Genome Canada and the Ontario Genomics Institute (OGI-051) at the Toronto Centre for Phenogenomics and obtained from the Canadian Mouse Mutant Repository.

## Funding information

This work was supported by a Wellcome Trust Investigator Award (215574/Z/19/Z) to DG, and a Biotechnology and Biological Sciences Research Council (BBSRC) grant BB/S014357/1 to CS.

## Data availability

Raw data for all nematode, *Drosophila* and mouse cohorts are provided.

## Supplementary Figures

**Figure S1.**
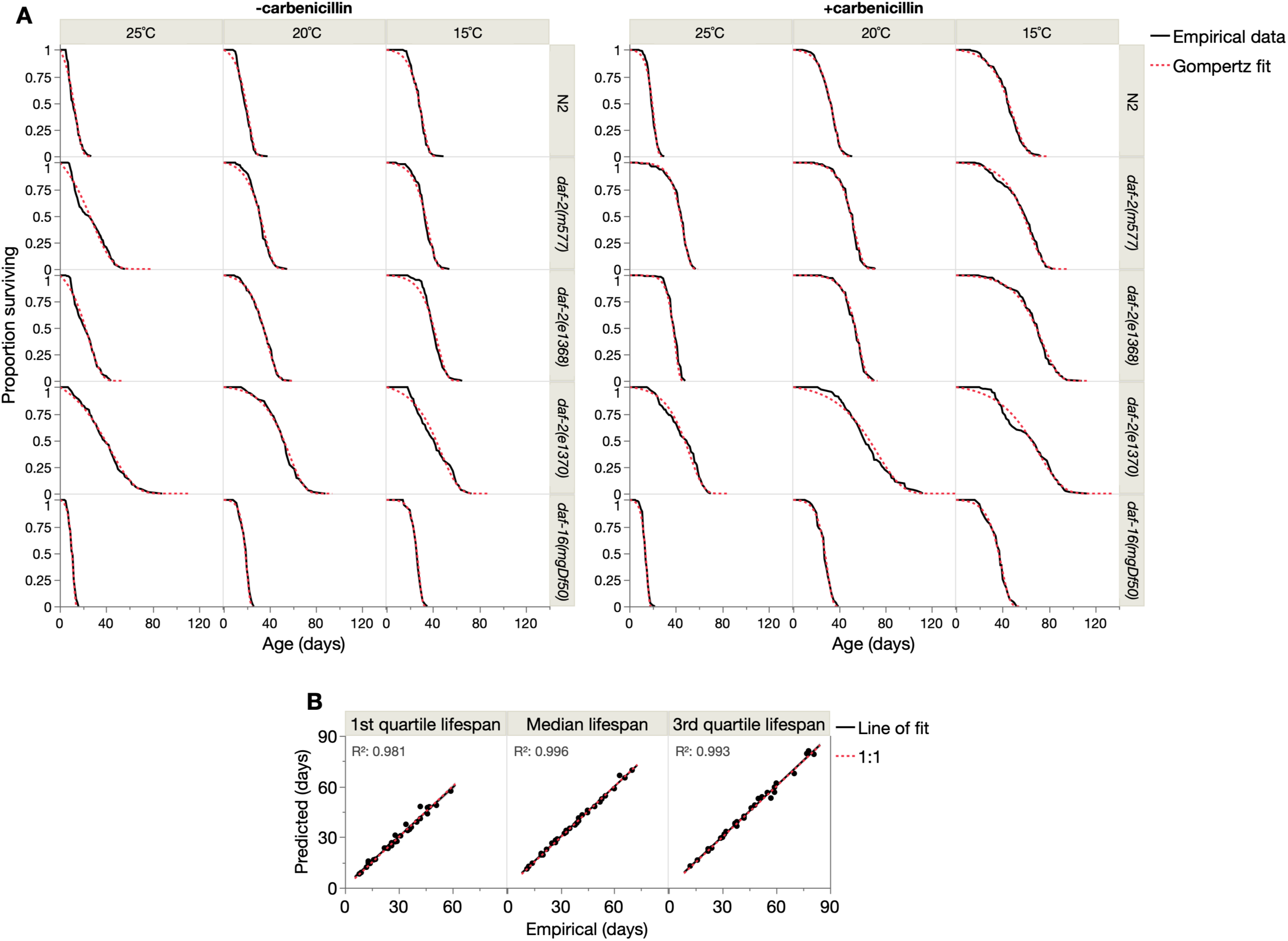
| The Gompertz model provides an appropriate fit to the 30 cohorts. (**A**) Overlay of empirical and idealized Gompertz curves for each of the 30 cohorts (left panel: cohorts without carbenicillin; right panel: carbenicillin-treated cohorts). (**B**) Linear regressions between true (empirical) and predicted (from the Gompertz fit) lifespan measures, across the 30 cohorts. First quartile, median and third quartile lifespans are the ages at which survival proportion equals, respectively, 0.75, 0.5 and 0.25; the results indicate strong prediction of overall survival curve shape (i.e. covering ages of early, middle and late deaths) by the Gompertz model for these cohorts. 95% confidence regions around the regression fits are shaded (barely visible); fits were assessed by F-tests.

**Figure S2.**
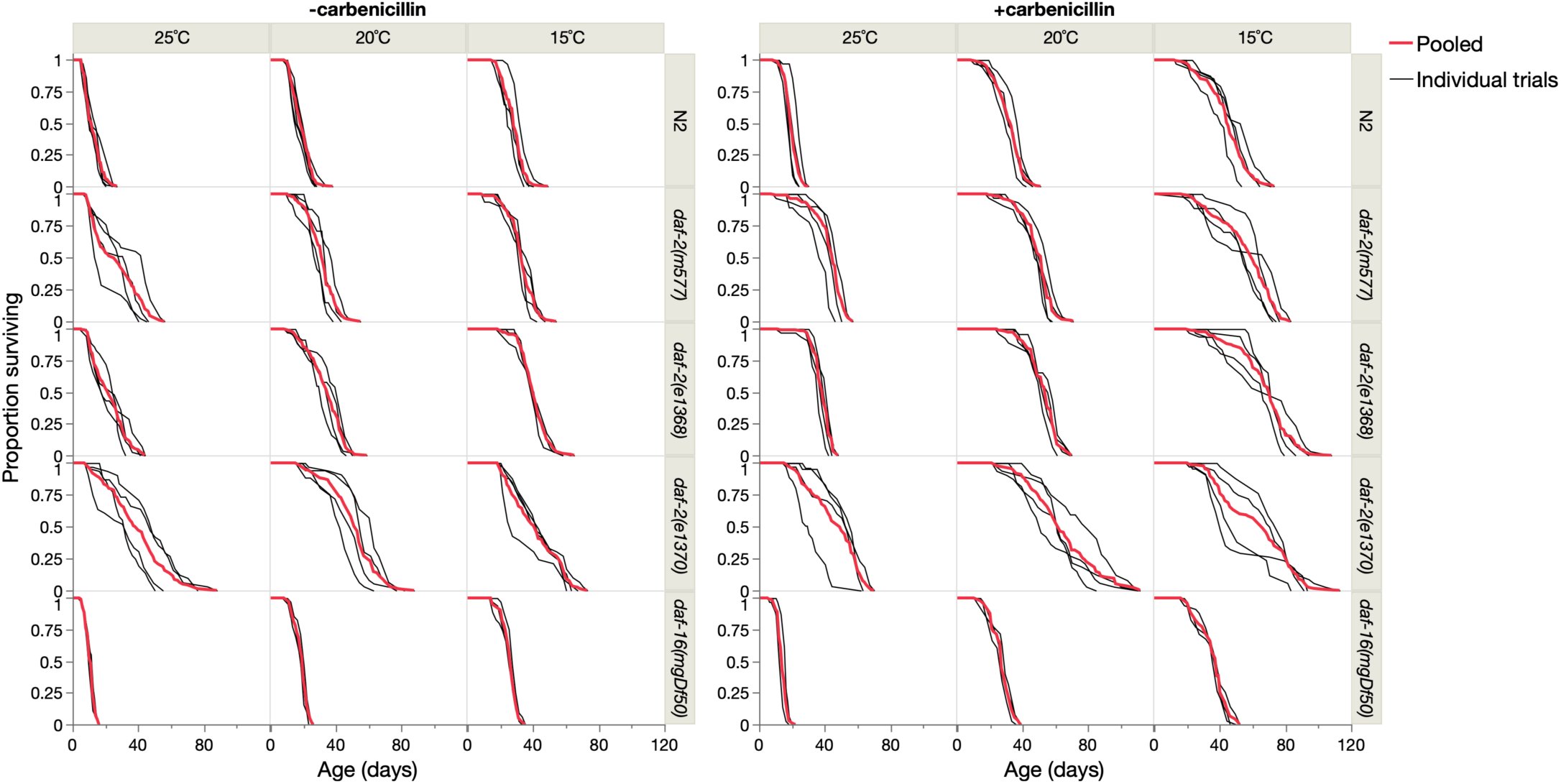
| Survival curves of the pooled and individual trial data for the 30 cohorts. Left panel: cohorts without carbenicillin; right panel: carbenicillin-treated cohorts. Survival proportions are from Kaplan-Meier survival analysis in JMP, with right censoring. All cohorts have at least 3 individual trials. Statistical details of the individual and pooled trials are provided in Table S1.

**Figure S3.**
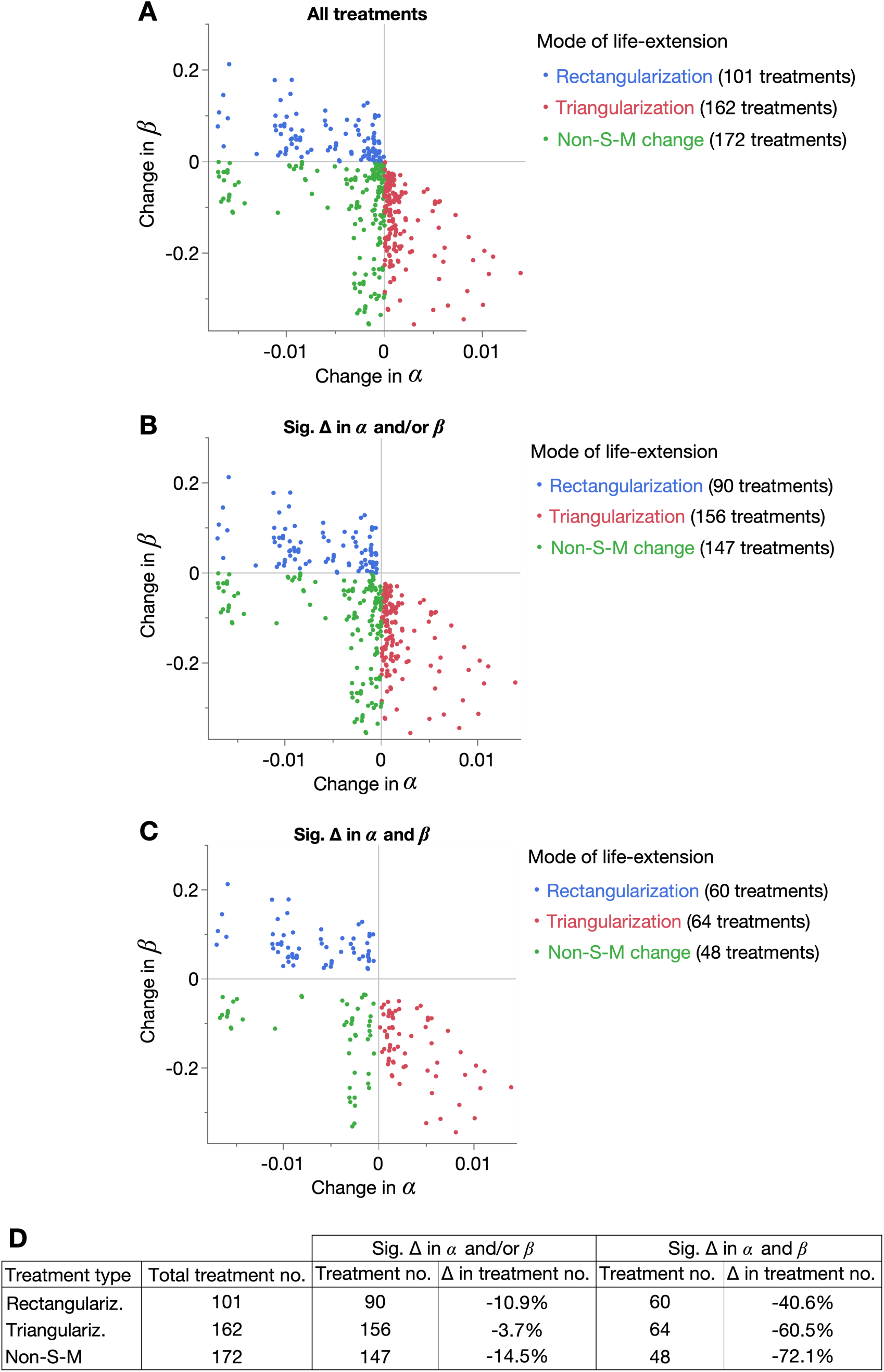
| Rectangularization and triangularization in the dataset are not statistical artifacts of fitting the Gompertz model. Changes in *β* plotted over the corresponding changes in *α*, for (**A**) all 435 pairwise treatments, (**B**) only treatments in which at least one of *α* and *β* are statistically significantly changed (likelihood ratio test, Benjamini-Hochberg corrected *p*<0.05), and (**C**) only treatments in which both *α* and *β* are statistically significantly changed. Change direction was always determined based on the longer-lived cohort minus shorter-lived cohort, such that all changes reflect the effects of life-extension. (**D**) Summary table of the number of treatments and percent change in this number for the conditions in **B** and **C**, relative to the total number of treatments.

**Figure S4.**
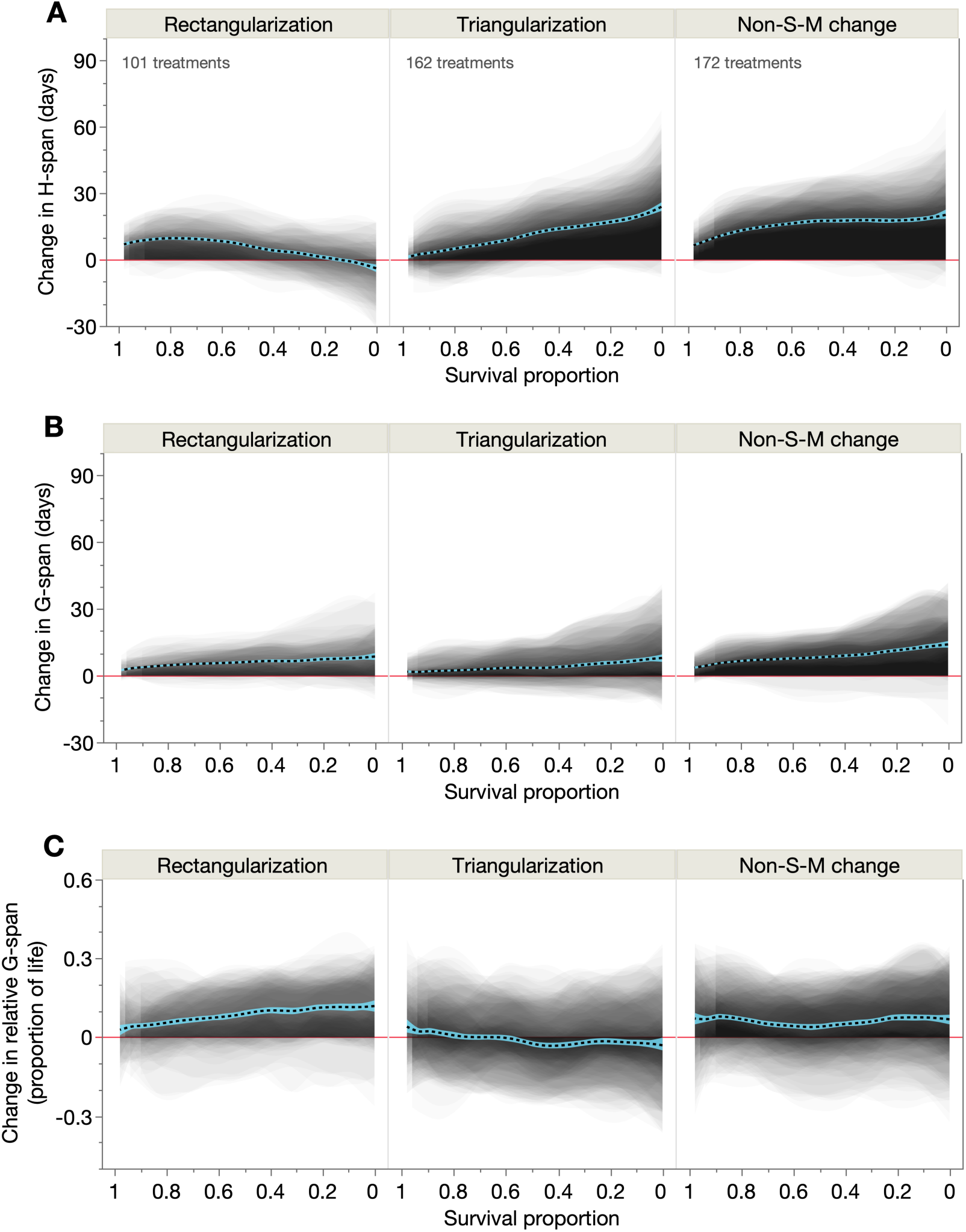
| Biodemographic determinants of rectangularization and triangularization are robust to the definition of H-span and G-span. Shaded and overlaid area plots for all treatments (comparison pairs) of each demographic treatment type (rectangularization, triangularization, non-S-M change), of the change in (**A**) H-span, (**B**) G-span, and (**C**) relative G-span, where H-span is the duration of locomotory capacity (sinusoidal or not) and G-span is the duration of end-of-life immotility (i.e. zero locomotory capacity). These shaded changes are plotted over survival proportion (x-axis left: shorter-lived population members, x-axis right: longer-lived population members). Red line: y=0, below which the lifespan-extending treatment shortens that trait. Dashed black line: summary spline smoother (lambda=0.05) of the 435 treatment changes, showing its 95% confidence region in blue. The number of treatments in each demographic type is annotated in **A**.

**Figure S5.**
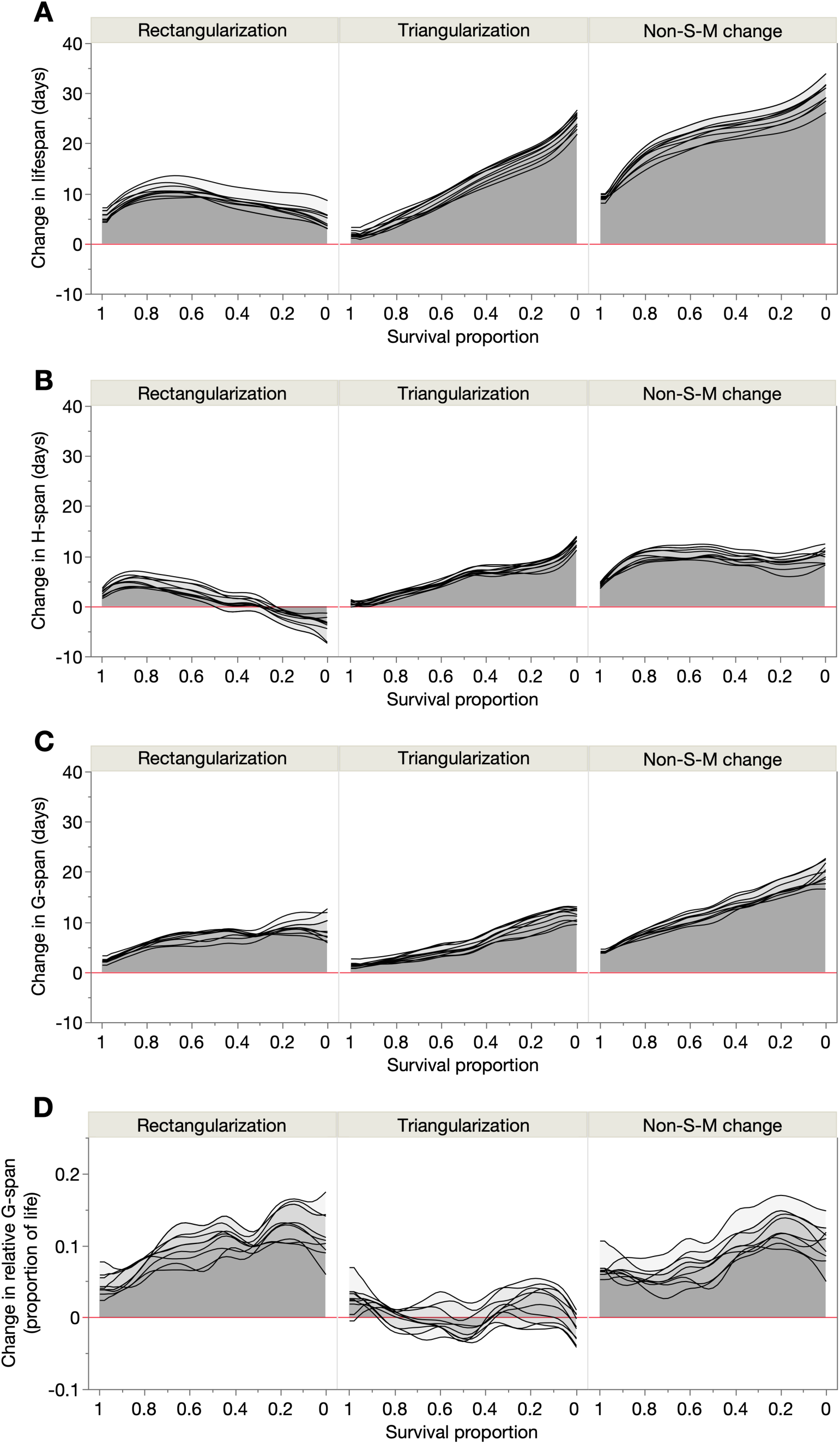
| Biodemographic determinants of rectangularization and triangularization are not artifacts of pairwise analysis. 10 summary spline smoothers (lambda=0.05) corresponding to 10 randomly subsampled pairwise treatments (from the 435 treatments) that cause rectangularization, triangularization or non-S-M changes, showing their effects on (**A**) lifespan, (**B**) H-span, (**C**) G-span, and (**D**) relative G-span. Each subsample contains the maximum possible number of treatments such that no cohort appears more than once: rectangularization (14 treatments), triangularization (15 treatments) and non-S-M changes (15 treatments). The area under each smoother is shaded, and they are plotted over survival proportion (x-axis left: shorter-lived population members, x-axis right: longer-lived population members). Red line: y=0, below which the lifespan-extending treatment shortens that trait.

**Figure S6.**
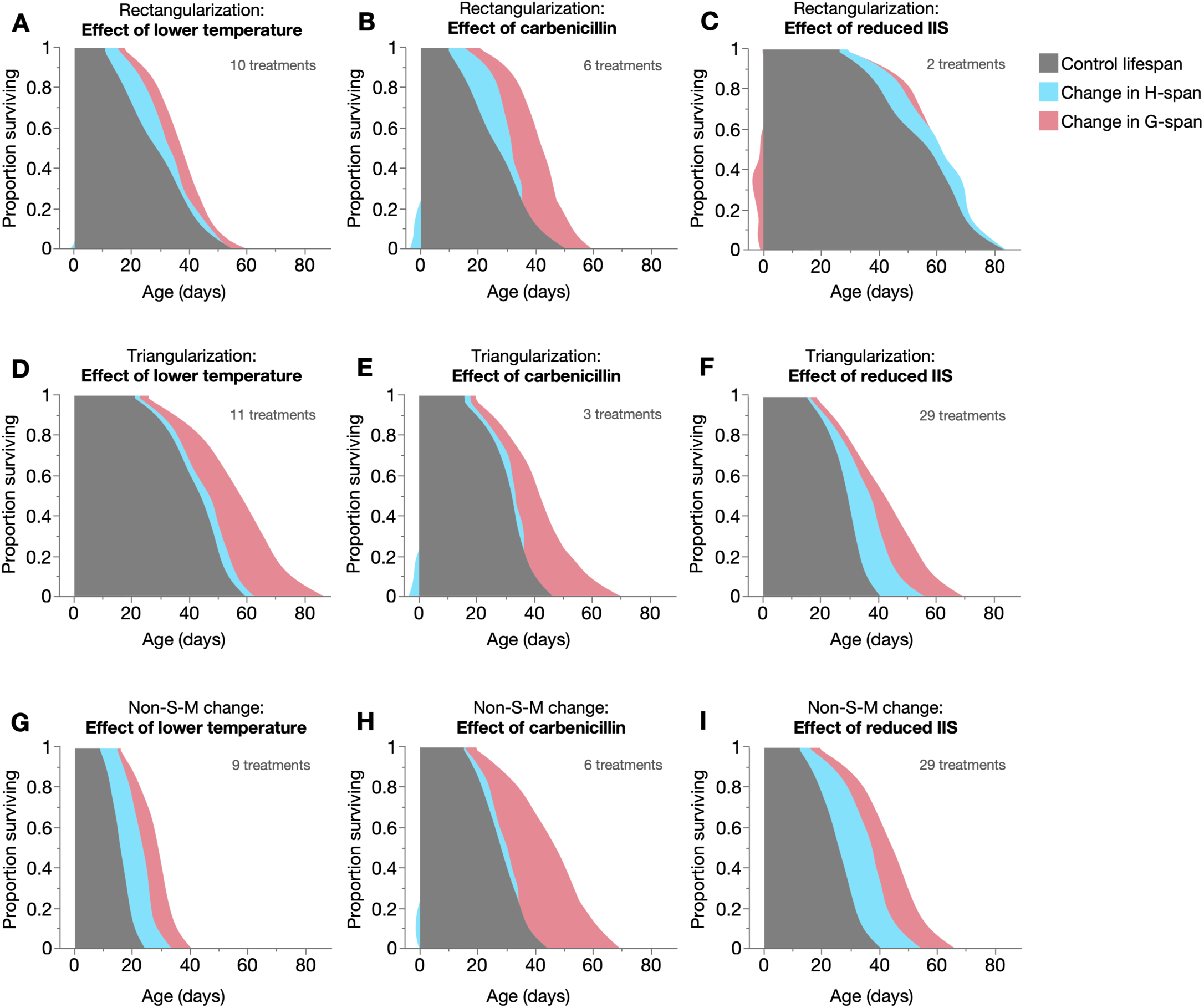
| Effects of specific interventions on H-span and G-span across demographic life-extension modes. Empirical summary figures of how lowered temperature (**A**, **D**, **G**), carbenicillin (**B**, **E**, **H**), and IIS modulation (**C**, **F**, **I**) cause rectangularizing (**A**–**C**), triangularizing (**D**–**F**) or non-S-M (**G**–**I**) effects on lifespan, through additive changes in H-span and G-span (spline smoother of each, stacked on spline smoother of control lifespan), displayed in a shaded survival curve-like format. Shortening of H-span or G-span is represented as a negative change relative to Age=0. These treatments include only those in which one condition (out of temperature, antibiotic and genotype) is changed at a time, thereby enabling the effects of each condition to be isolated; the numbers of these treatments are annotated. Note that the number of such temperature-only, antibiotic-only and IIS-only treatments is set by the number of ‘levels’ within each conditions (3 temperatures, ± carbenicillin, 5 genotypes), which is 30 temperature-only treatments, 15 antibiotic-only and 60 IIS-only treatments (which are here distributed between the 3 demographic life-extension modes).

**Figure S7.**
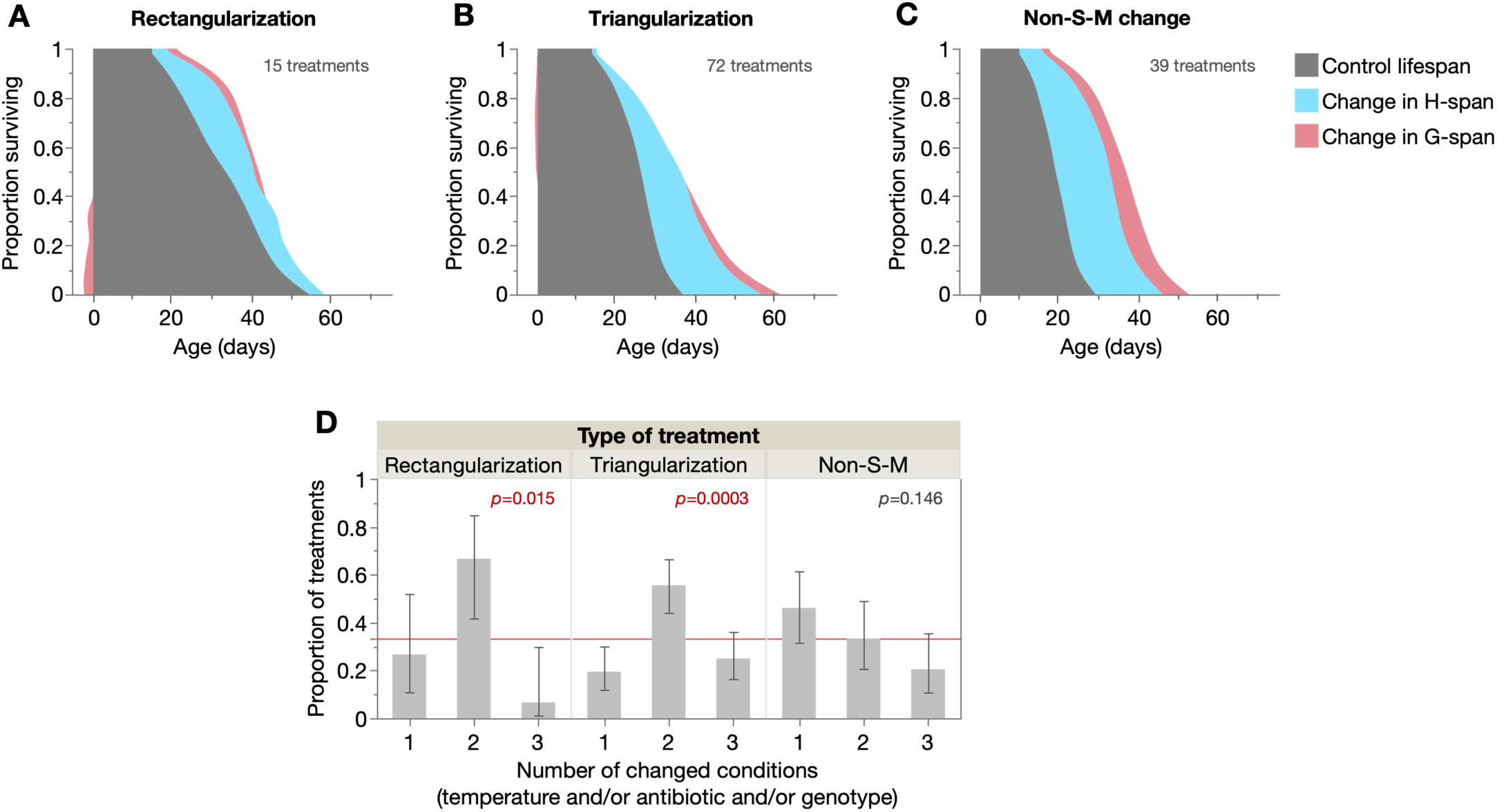
| Different biodemographic modes of morbidity compression. (**A**–**C**) Empirical summary figures of how rectangularizing, triangularizing and non-S-M treatments that compress morbidity (decrease relative G-span) extend lifespan through additive changes in H-span and G-span (spline smoother of each, stacked on spline smoother of control lifespan), displayed in a shaded survival curve-like format. Mild shortening of G-span in the longest-lived population members in rectangularizing treatments and shorter-lived population members in triangularizing treatments is represented as negative changes relative to Age=0. (**D**) Proportions of the relative G-span-reducing rectangularizing, triangularizing and non-S-M treatments that arise from one, two or three changes in experiment condition (temperature, ± carbenicillin, genotype). Proportions within each panel sum to 1, and proportions above or below the red line (expected proportion under null hypothesis: 0.33) indicate enrichment or depletion of that number of changed conditions. 95% confidence intervals are shown and Pearson chi-square goodness-of-fit tests were run for each panel (*p* values annotated), with sample sizes of, rectangularizing: 15 treatments, triangularizing: 72 treatments, and non-S-M: 39 treatments.

**Figure S8.**
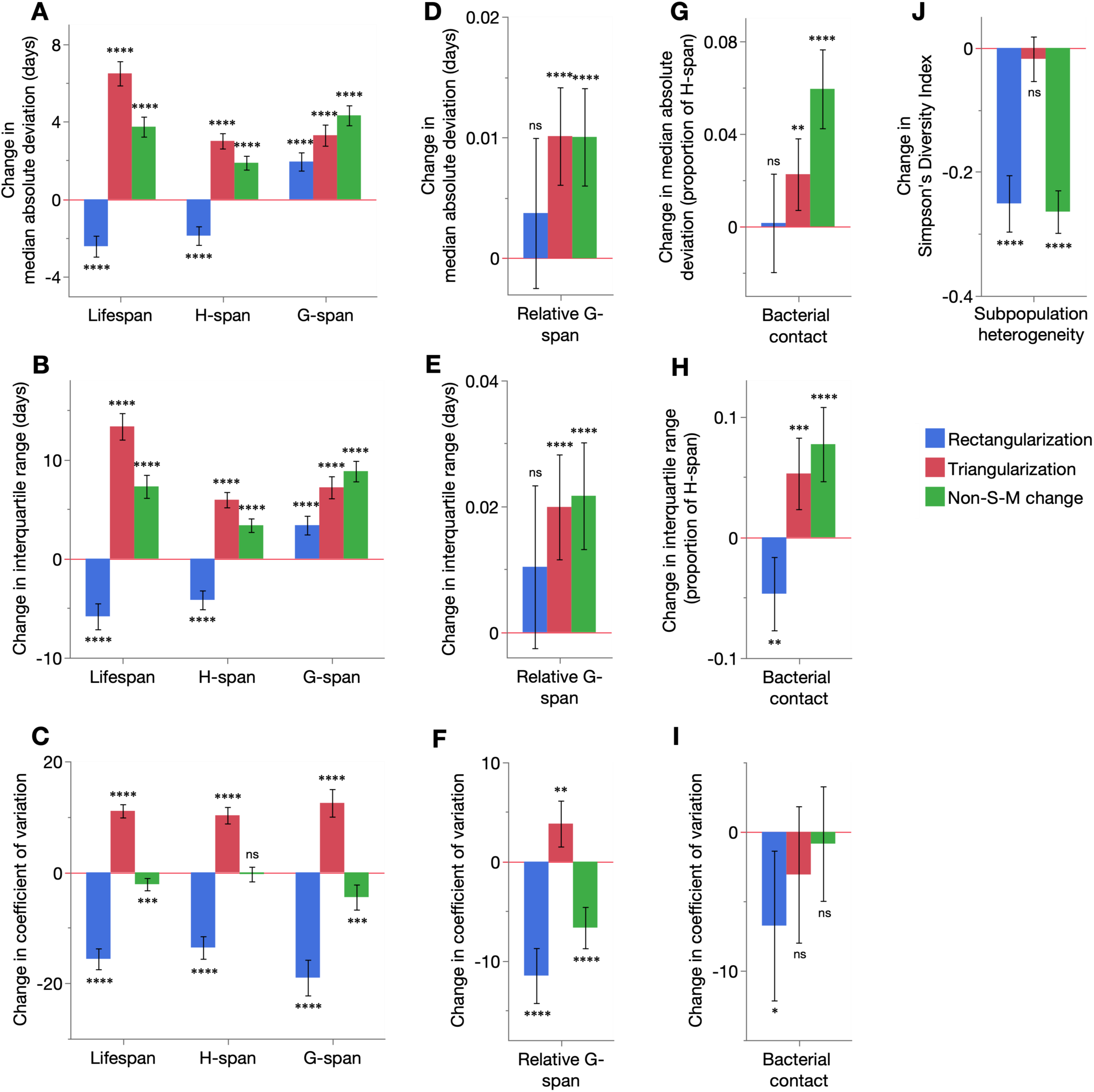
| Effects of different demographic life-extension modes on variation in the aging process. Mean changes in the (**A**, **D**, **G**) median absolute deviation, (**B**, **E**, **H**) interquartile range and (**C**, **F**, **I**) coefficient of variation of (**A**–**C**) lifespan, H-span and G-span, (**D**–**F**) relative G-span and (**G**–**I**) bacterial contact, by rectangularizing, triangularizing and non-S-M treatments. (**J**) Mean changes in population heterogeneity resulting from changes in subpopulation (PIC, pIC, pnIC) proportions by rectangularizing, triangularizing and non-S-M treatments, as measured by Simpson’s Diversity Index. Mean changes in all panels were assessed with two-tailed one sample t-tests (*H*_0_: mean change=0), showing 95% confidence intervals; Benjamini-Hochberg correction of *p* values did not change which treatments reached the 0.05 significance threshold (data not shown). ns *p* > 0.05, * *p* ≤ 0.05, ** *p* ≤ 0.01, *** *p* ≤ 0.001, **** *p* ≤ 0.0001.

**Figure S9.**
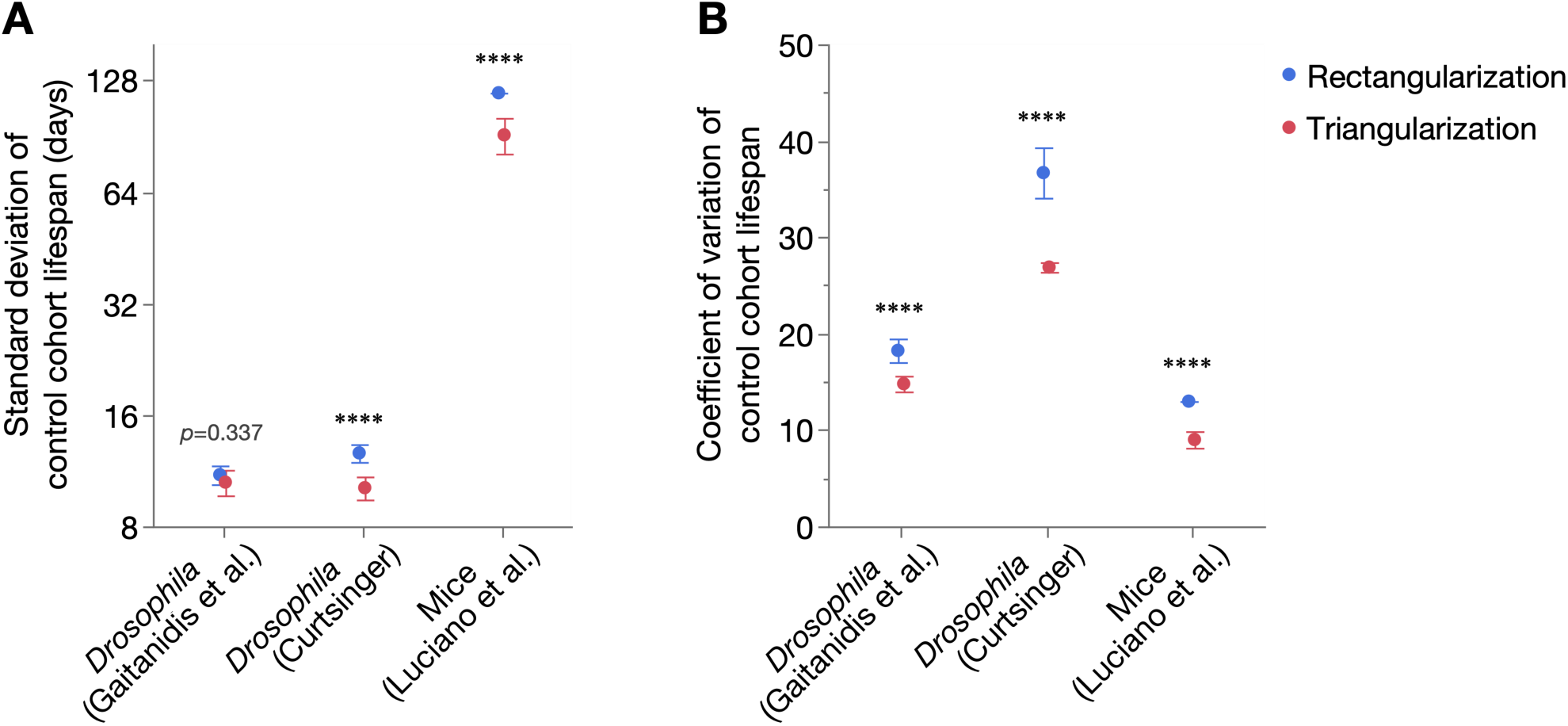
| Greater lifespan variation in cohorts undergoing survival curve rectangularization than triangularization. Mean (**A**) standard deviation and (**B**) coefficient of variation of lifespan of control cohorts that underwent rectangularizing versus triangularizing treatments (respectively, for Gaitanidis et al. (2019), Curtsinger (2015), Luciano et al. (2024): *n*=37, 33; 15, 10; 3, 15). Differences in means were assessed with two-tailed Student’s t-tests, showing 95% confidence intervals. In all panels: ns *p* > 0.05, * *p* ≤ 0.05, ** *p* ≤ 0.01, *** *p* ≤ 0.001, **** *p* ≤ 0.0001.

**Figure S10.**
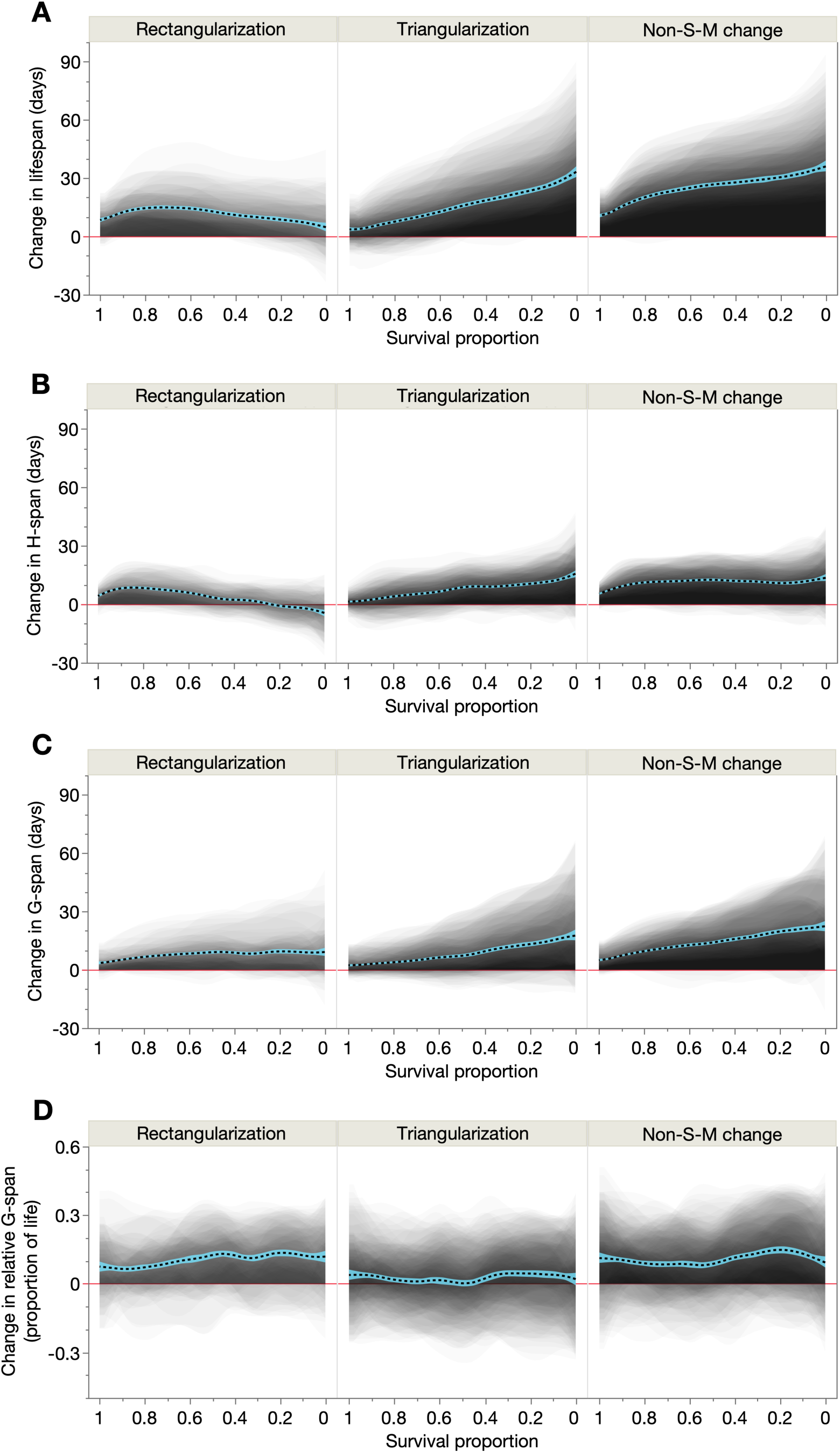
| Biodemographic determinants of rectangularization and triangularization are unaffected by the exclusion of three cohorts. Shaded and overlaid area plots for all treatments (except those involving 3 cohorts that did not pass Anderson-Darling tests for goodness-of-fit to Gompertz: N2 15°C, *daf-2(m577)* 15°C, *daf-2(m577)* 20°C carb.; see Table S2) of each demographic treatment type (rectangularization, triangularization, non-S-M change), of the change in (**A**) lifespan, (**B**) H-span, (**C**) G-span and (**D**) relative G-span. These shaded changes are plotted over survival proportion (x-axis left: shorter-lived population members, x-axis right: longer-lived population members). Red line: y=0, below which the lifespan-extending treatment shortens that trait. Dashed black line: summary spline smoother (lambda=0.05) of the 435 treatment changes, showing its 95% confidence region in blue.

## Supplementary Tables

**Table S1.** | Lifespan and Gompertz statistics for all cohorts and trials. N2, wild type. [C], combined (pooled) data from all trials; carb., carbenicillin. Mean lifespan and its standard errors were obtained by Kaplan-Meier survival analysis in JMP, and Gompertz parameters and their associated statistics (LCI: lower 95% confidence interval; UCI: upper 95% confidence interval) obtained by maximum likelihood estimation in WinModest. In each cohort section, each row below the first corresponds to one trial. Although experiments for the 30 cohorts were not always performed simultaneously due to practical constraints, inter-trial lifespan variation was well controlled (Fig. S2); such variation was greater in the *daf-2* mutants, a property of these strains noted in previous studies from this laboratory. The analyses in this study utilize the pooled data [C] from all trials, so lifespan and Gompertz statistics are not displayed for individual trials. Two additional larger trials were performed for all N2, *daf-2(m577)*, *daf-2(e1368)* and *daf-2(e1370)* cohorts, whose raw data is publicly available (4).

| Cohort | No. dead/<br>censors | Mean lifespan<br>(days since L4) | Standard error<br>of lifespan | $\alpha$ | $\beta$ | LCI($\alpha$) | UCI($\alpha$) | LCI($\beta$) | UCI($\beta$) | Negative log-likelihood |
| --- | --- | --- | --- | --- | --- | --- | --- | --- | --- | --- |
| N2 25 °C | [C] 169/8<br>70/1<br>33/2<br>31/4<br>35/1 | 12.6<br>11.3<br>13.7<br>11.6<br>14.9 | 0.38 | 0.01707 | 0.15825 | 0.01212 | 0.02405 | 0.13550 | 0.18483 | -523.62398 |
| N2 20 °C | [C] 246/14<br>67/2<br>33/3<br>33/3<br>36/0<br>44/2<br>33/4 | 19.0<br>19.9<br>20.6<br>18.1<br>17.4<br>19.8<br>16.8 | 0.34 | 0.00488 | 0.16431 | 0.00347 | 0.00686 | 0.14833 | 0.18202 | -792.21807 |
| N2 15 °C | [C] 159/18<br>62/8<br>33/3<br>30/5<br>34/2 | 28.3<br>27.4<br>33.5<br>25.2<br>27.3 | 0.47 | 0.00124 | 0.15477 | 0.00074 | 0.00210 | 0.13816 | 0.17337 | -529.86476 |
| N2 25 °C carb. | [C] 173/6<br>70/2<br>34/1<br>35/1<br>34/2 | 19.8<br>18.2<br>20.8<br>24.0<br>17.6 | 0.30 | 0.00104 | 0.25251 | 0.00057 | 0.00188 | 0.22520 | 0.28313 | -497.43075 |
| N2 20 °C carb. | [C] 173/6<br>70/1<br>33/3<br>35/1<br>35/1 | 31.8<br>29.0<br>32.2<br>32.4<br>36.4 | 0.59 | 0.00101 | 0.13889 | 0.00058 | 0.00176 | 0.12314 | 0.15666 | -604.25161 |
| N2 15 °C carb. | [C] 161/10<br>62/1<br>32/4<br>34/2<br>33/3 | 44.3<br>38.2<br>50.2<br>47.0<br>47.3 | 0.96 | 0.00120 | 0.08546 | 0.00072 | 0.00199 | 0.07523 | 0.09709 | -634.88993 |
| daf-2(m577) 25 °C | [C] 125/13<br>21/0<br>32/2<br>31/5<br>41/6 | 26.3<br>26.6<br>34.7<br>26.8<br>19.1 | 1.20 | 0.01125 | 0.05703 | 0.00755 | 0.01677 | 0.04502 | 0.07225 | -502.48843 |
| daf-2(m577) 20 °C | [C] 119/22<br>32/4<br>26/7<br>29/7<br>32/4 | 31.3<br>36.0<br>33.2<br>28.3<br>27.7 | 0.76 | 0.00215 | 0.11245 | 0.00128 | 0.00360 | 0.09787 | 0.12920 | -437.08262 |
| daf-2(m577) 15 °C | [C] 136/5<br>35/1<br>30/3<br>35/1<br>36/0 | 33.0<br>34.2<br>33.4<br>32.7<br>31.8 | 0.65 | 0.00100 | 0.13372 | 0.00055 | 0.00182 | 0.11744 | 0.15227 | -476.21816 |
| daf-2(m577) 25 °C carb. | [C] 104/12<br>18/1<br>29/2<br>22/8<br>35/1 | 43.2<br>35.8<br>45.8<br>41.9<br>45.6 | 0.84 | 0.00010 | 0.15699 | 0.00003 | 0.00029 | 0.13438 | 0.18341 | -359.32556 |
| daf-2(m577) 20 °C carb. | [C] 123/17<br>30/2<br>27/9<br>33/3<br>33/3 | 49.3<br>47.5<br>53.7<br>46.4<br>50.3 | 0.81 | 0.00015 | 0.12508 | 0.00007 | 0.00035 | 0.10962 | 0.14272 | -446.00468 |
| daf-2(m577) 15 °C carb. | [C] 129/15<br>32/4<br>30/6<br>34/2<br>33/3 | 56.2<br>51.7<br>56.9<br>63.6<br>52.8 | 1.38 | 0.00059 | 0.07684 | 0.00030 | 0.00115 | 0.06617 | 0.08923 | -534.37366 |
| daf-2(e1368) 25 °C | [C] 141/15<br>35/1<br>34/2<br>30/6<br>42/6 | 22.3<br>23.2<br>23.3<br>22.8<br>20.7 | 0.80 | 0.00964 | 0.08711 | 0.00657 | 0.01415 | 0.07272 | 0.10435 | -526.02146 |
| daf-2(e1368) 20 °C | [C] 118/24<br>31/5<br>26/8<br>32/4<br>29/7 | 33.9<br>34.6<br>37.5<br>30.8<br>33.2 | 0.90 | 0.00177 | 0.10677 | 0.00101 | 0.00311 | 0.09190 | 0.12404 | -445.63139 |
| daf-2(e1368) 15 °C | [C] 132/12<br>34/2<br>33/3<br>34/2<br>31/5 | 40.5<br>39.3<br>41.0<br>41.5<br>40.1 | 0.69 | 0.00065 | 0.11689 | 0.00035 | 0.00118 | 0.10338 | 0.13216 | -477.79654 |
| *daf-2(e1368)* 25 °C carb. | [C] 127/17<br>33/3<br>32/4<br>34/2<br>28/8 | 38.4<br>35.6<br>37.1<br>40.4<br>40.2 | 0.45 | 0.00002 | 0.23469 | 0.00000 | 0.00006 | 0.20498 | 0.26872 | -389.60542 |
| *daf-2(e1368)* 20 °C carb. | [C] 104/32<br>25/5<br>19/15<br>29/7<br>31/5 | 52.8<br>52.3<br>56.3<br>50.6<br>53.1 | 0.83 | 0.00006 | 0.13500 | 0.00002 | 0.00018 | 0.11674 | 0.15612 | -373.32477 |
| *daf-2(e1368)* 15 °C carb. | [C] 123/21<br>30/6<br>30/6<br>35/1<br>28/8 | 67.4<br>60.6<br>66.8<br>70.3<br>71.1 | 1.42 | 0.00037 | 0.07013 | 0.00019 | 0.00072 | 0.06115 | 0.08043 | -517.81885 |
| *daf-2(e1370)* 25 °C | [C] 143/10<br>36/0<br>31/2<br>34/2<br>42/6 | 39.2<br>32.2<br>48.6<br>29.3<br>46.3 | 1.44 | 0.00616 | 0.04593 | 0.00427 | 0.00888 | 0.03827 | 0.05512 | -616.37967 |
| *daf-2(e1370)* 20 °C | [C] 130/12<br>33/1<br>30/6<br>31/5<br>36/0 | 50.9<br>55.8<br>58.8<br>41.6<br>47.8 | 1.22 | 0.00111 | 0.07299 | 0.00065 | 0.00191 | 0.06352 | 0.08387 | -533.20004 |
| *daf-2(e1370)* 15 °C | [C] 126/18<br>30/6<br>34/2<br>33/3<br>29/7 | 41.5<br>44.3<br>34.9<br>43.1<br>44.2 | 1.28 | 0.00276 | 0.06663 | 0.00170 | 0.00448 | 0.05648 | 0.07861 | -525.3664 |
| *daf-2(e1370)* 25 °C carb. | [C] 112/36<br>32/4<br>22/8<br>30/6<br>28/18 | 46.2<br>31.1<br>53.3<br>48.0<br>52.9 | 1.30 | 0.00118 | 0.08054 | 0.00064 | 0.00218 | 0.06862 | 0.09453 | -462.67365 |
| *daf-2(e1370)* 20 °C carb. | [C] 126/17<br>33/3<br>28/7<br>33/3<br>32/4 | 64.5<br>62.7<br>76.1<br>58.1<br>61.5 | 1.77 | 0.00156 | 0.04608 | 0.00097 | 0.00253 | 0.03957 | 0.05367 | -571.01596 |
| *daf-2(e1370)* 15 °C carb. | [C] 124/9<br>20/5<br>34/2<br>34/2<br>36/0 | 62.6<br>51.6<br>53.3<br>73.4<br>67.4 | 1.90 | 0.00151 | 0.04821 | 0.00090 | 0.00254 | 0.04097 | 0.05673 | -556.58801 |
| *daf-16(mgDf50)* 25 °C | [C] 102/5<br>36/0<br>33/1<br>33/4 | 10.8<br>10.3<br>11.1<br>11.0 | 0.26 | 0.00316 | 0.40240 | 0.00156 | 0.00642 | 0.34399 | 0.47073 | -247.34718 |
| *daf-16(mgDf50)* 20 °C | [C] 103/4<br>36/0<br>35/0<br>32/4 | 18.8<br>18.1<br>20.2<br>18.2 | 0.38 | 0.00058 | 0.30324 | 0.00023 | 0.00142 | 0.25986 | 0.35386 | -283.16956 |
| *daf-16(mgDf50)* 15 °C | [C] 100/6<br>35/1<br>33/1<br>32/4 | 26.1<br>24.8<br>25.7<br>27.9 | 0.44 | 0.00015 | 0.26535 | 0.00005 | 0.00044 | 0.22802 | 0.30879 | -289.4465 |
| *daf-16(mgDf50)* 25 °C carb. | [C] 99/8<br>36/0<br>32/4<br>31/4 | 14.2<br>13.0<br>14.0<br>15.7 | 0.27 | 0.00118 | 0.37083 | 0.00054 | 0.00259 | 0.32115 | 0.42820 | -246.25844 |
| *daf-16(mgDf50)* 20 °C carb. | [C] 90/8<br>35/1<br>25/4<br>30/3 | 27.3<br>26.3<br>28.3<br>27.5 | 0.60 | 0.00062 | 0.19110 | 0.00026 | 0.00148 | 0.16240 | 0.22486 | -286.83034 |
| *daf-16(mgDf50)* 15 °C carb. | [C] 94/11<br>35/1<br>34/2<br>25/8 | 36.0<br>37.1<br>35.6<br>35.1 | 0.85 | 0.00062 | 0.13495 | 0.00028 | 0.00137 | 0.11494 | 0.15845 | -334.93058 |

**Table S2.**
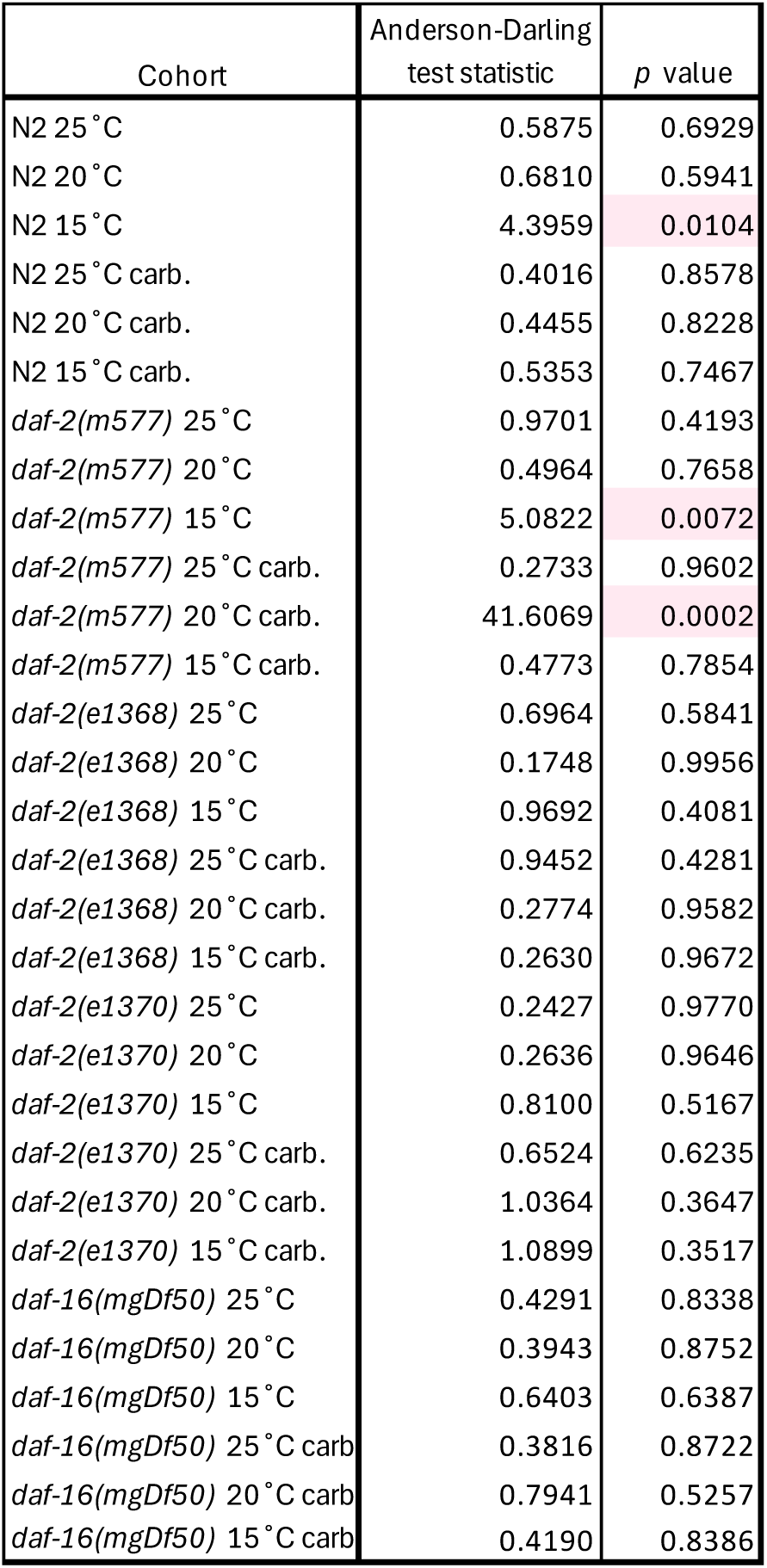
| Anderson-Darling goodness-of-fit test statistics for all cohorts. Test statistics and *p* values for parametric bootstrap Anderson-Darling tests, to assess goodness-of-fit of the 30 cohorts to the MLE Gompertz distributions. This test assesses overall, tail-weighted fit (here as the sum of squared, tail-weighted deviations between these functions across all ages). For each cohort, observed A-D statistics were compared against null distributions generated from 5000 parametric bootstrap populations simulated in JMP under the fitted Gompertz distribution of interest (MLE parameters obtained in WinModest), by generating random survival proportions (Uniform(0,1)) and population sizes matching the observed populations (including censored individuals). Monte Carlo *p* values were computed as the proportion of bootstrap A-D statistics greater than or equal to the observed statistic, applying the Phipson–Smyth correction (addition of 1 to numerator and denominator). Three *p* values less than 0.05 are shaded. Exclusion of these three cohorts does not alter the overall dynamics of healthspan and gerospan in rectangularization, triangularization and non-S-M changes (Fig. S10).

| Cohort | Anderson-Darling<br>test statistic | *p* value |
| --- | --- | --- |
| N2 25 °C | 0.5875 | 0.6929 |
| N2 20 °C | 0.6810 | 0.5941 |
| N2 15 °C | 4.3959 | 0.0104 |
| N2 25 °C carb. | 0.4016 | 0.8578 |
| N2 20 °C carb. | 0.4455 | 0.8228 |
| N2 15 °C carb. | 0.5353 | 0.7467 |
| *daf-2(m577)* 25 °C | 0.9701 | 0.4193 |
| *daf-2(m577)* 20 °C | 0.4964 | 0.7658 |
| *daf-2(m577)* 15 °C | 5.0822 | 0.0072 |
| *daf-2(m577)* 25 °C carb. | 0.2733 | 0.9602 |
| *daf-2(m577)* 20 °C carb. | 41.6069 | 0.0002 |
| *daf-2(m577)* 15 °C carb. | 0.4773 | 0.7854 |
| *daf-2(e1368)* 25 °C | 0.6964 | 0.5841 |
| *daf-2(e1368)* 20 °C | 0.1748 | 0.9956 |
| *daf-2(e1368)* 15 °C | 0.9692 | 0.4081 |
| *daf-2(e1368)* 25 °C carb. | 0.9452 | 0.4281 |
| *daf-2(e1368)* 20 °C carb. | 0.2774 | 0.9582 |
| *daf-2(e1368)* 15 °C carb. | 0.2630 | 0.9672 |
| *daf-2(e1370)* 25 °C | 0.2427 | 0.9770 |
| *daf-2(e1370)* 20 °C | 0.2636 | 0.9646 |
| *daf-2(e1370)* 15 °C | 0.8100 | 0.5167 |
| *daf-2(e1370)* 25 °C carb. | 0.6524 | 0.6235 |
| *daf-2(e1370)* 20 °C carb. | 1.0364 | 0.3647 |
| *daf-2(e1370)* 15 °C carb. | 1.0899 | 0.3517 |
| *daf-16(mgDf50)* 25 °C | 0.4291 | 0.8338 |
| *daf-16(mgDf50)* 20 °C | 0.3943 | 0.8752 |
| *daf-16(mgDf50)* 15 °C | 0.6403 | 0.6387 |
| *daf-16(mgDf50)* 25 °C carb | 0.3816 | 0.8722 |
| *daf-16(mgDf50)* 20 °C carb | 0.7941 | 0.5257 |
| *daf-16(mgDf50)* 15 °C carb | 0.4190 | 0.8386 |

**Table S3.** | Morbidity-compressing rectangularizing treatments. Temperature, antibiotic usage and genotype of rectangularizing treatments that decreased relative G-span. Each treatment comprises a pair of cohorts (control and treatment cohorts), and each row contains one treatment (15 in total).

| Rectangularising S-M treatments |  |  |  |  |  |
| --- | --- | --- | --- | --- | --- |
| Control cohort |  |  | Treatment cohort |  |  |
| Temperature | ± Carbenicillin | Genotype | Temperature | ± Carbenicillin | Genotype |
| 25 °C | Carb | N2 | 15 °C | No carb | *daf-16(mgDf50)* |
| 25 °C | No carb | *daf-2(m577)* | 15 °C | No carb | *daf-2(e1368)* |
| 15 °C | Carb | *daf-2(e1370)* | 15 °C | Carb | *daf-2(e1368)* |
| 25 °C | No carb | *daf-2(m577)* | 20 °C | No carb | *daf-2(e1368)* |
| 15 °C | Carb | *daf-16(mgDf50)* | 20 °C | Carb | *daf-2(e1368)* |
| 20 °C | Carb | *daf-2(m577)* | 20 °C | Carb | *daf-2(e1368)* |
| 15 °C | Carb | N2 | 20 °C | Carb | *daf-2(e1368)* |
| 15 °C | Carb | *daf-16(mgDf50)* | 25 °C | Carb | *daf-2(e1368)* |
| 20 °C | Carb | *daf-16(mgDf50)* | 25 °C | Carb | *daf-2(e1368)* |
| 20 °C | Carb | N2 | 25 °C | Carb | *daf-2(e1368)* |
| 25 °C | No carb | *daf-2(m577)* | 15 °C | No carb | *daf-2(m577)* |
| 25 °C | No carb | *daf-2(m577)* | 20 °C | No carb | *daf-2(m577)* |
| 15 °C | Carb | *daf-16(mgDf50)* | 25 °C | Carb | *daf-2(m577)* |
| 20 °C | Carb | N2 | 25 °C | Carb | *daf-2(m577)* |
| 25 °C | No carb | *daf-2(m577)* | 15 °C | No carb | N2 |

**Table S4.** | Morbidity-compressing triangularizing treatments. Temperature, antibiotic usage and genotype of triangularizing treatments that decreased relative G-span. Each treatment comprises a pair of cohorts (control and treatment cohorts), and each row contains one treatment (72 in total).

| Triangularising S-M treatments |  |  |  |  |  |
| --- | --- | --- | --- | --- | --- |
| Control cohort |  |  | Treatment cohort |  |  |
| Temperature | ± Carbenicillin | Genotype | Temperature | ± Carbenicillin | Genotype |
| 15°C | No carb | *daf-16(mgDf50)* | 15°C | No carb | *daf-2(e1368)* |
| 15°C | Carb | *daf-16(mgDf50)* | 15°C | No carb | *daf-2(e1368)* |
| 20°C | No carb | *daf-16(mgDf50)* | 15°C | No carb | *daf-2(e1368)* |
| 20°C | Carb | *daf-16(mgDf50)* | 15°C | No carb | *daf-2(e1368)* |
| 25°C | Carb | *daf-2(e1368)* | 15°C | No carb | *daf-2(e1368)* |
| 15°C | No carb | *daf-16(mgDf50)* | 20°C | No carb | *daf-2(e1368)* |
| 20°C | No carb | *daf-16(mgDf50)* | 20°C | No carb | *daf-2(e1368)* |
| 20°C | Carb | *daf-16(mgDf50)* | 20°C | No carb | *daf-2(e1368)* |
| 25°C | Carb | *daf-16(mgDf50)* | 20°C | No carb | *daf-2(e1368)* |
| 15°C | No carb | *daf-2(m577)* | 20°C | No carb | *daf-2(e1368)* |
| 15°C | No carb | N2 | 20°C | No carb | *daf-2(e1368)* |
| 20°C | Carb | N2 | 20°C | No carb | *daf-2(e1368)* |
| 25°C | Carb | N2 | 20°C | No carb | *daf-2(e1368)* |
| 20°C | No carb | *daf-16(mgDf50)* | 25°C | No carb | *daf-2(e1368)* |
| 25°C | No carb | *daf-16(mgDf50)* | 25°C | No carb | *daf-2(e1368)* |
| 25°C | Carb | *daf-16(mgDf50)* | 25°C | No carb | *daf-2(e1368)* |
| 20°C | No carb | N2 | 25°C | No carb | *daf-2(e1368)* |
| 25°C | Carb | N2 | 25°C | No carb | *daf-2(e1368)* |
| 15°C | Carb | *daf-16(mgDf50)* | 15°C | No carb | *daf-2(e1370)* |
| 20°C | Carb | *daf-16(mgDf50)* | 15°C | No carb | *daf-2(e1370)* |
| 25°C | Carb | *daf-16(mgDf50)* | 15°C | No carb | *daf-2(e1370)* |
| 20°C | Carb | N2 | 15°C | No carb | *daf-2(e1370)* |
| 25°C | Carb | N2 | 15°C | No carb | *daf-2(e1370)* |
| 15°C | Carb | *daf-16(mgDf50)* | 20°C | No carb | *daf-2(e1370)* |
| 20°C | Carb | *daf-16(mgDf50)* | 20°C | No carb | *daf-2(e1370)* |
| 20°C | Carb | *daf-2(m577)* | 20°C | No carb | *daf-2(e1370)* |
| 25°C | Carb | *daf-2(m577)* | 20°C | No carb | *daf-2(e1370)* |
| 20°C | Carb | N2 | 20°C | No carb | *daf-2(e1370)* |
| 15°C | Carb | *daf-2(e1370)* | 20°C | Carb | *daf-2(e1370)* |
| 15°C | Carb | *daf-2(m577)* | 20°C | Carb | *daf-2(e1370)* |
| 20°C | Carb | *daf-2(m577)* | 20°C | Carb | *daf-2(e1370)* |
| 15°C | No carb | *daf-16(mgDf50)* | 25°C | No carb | *daf-2(e1370)* |
| 15°C | Carb | *daf-16(mgDf50)* | 25°C | No carb | *daf-2(e1370)* |
| 20°C | No carb | *daf-16(mgDf50)* | 25°C | No carb | *daf-2(e1370)* |
| 20°C | Carb | *daf-16(mgDf50)* | 25°C | No carb | *daf-2(e1370)* |
| 25°C | No carb | *daf-16(mgDf50)* | 25°C | No carb | *daf-2(e1370)* |
| 25°C | Carb | *daf-16(mgDf50)* | 25°C | No carb | *daf-2(e1370)* |
| 25°C | Carb | *daf-2(e1368)* | 25°C | No carb | *daf-2(e1370)* |
| 15°C | No carb | *daf-2(m577)* | 25°C | No carb | *daf-2(e1370)* |
| 20°C | No carb | *daf-2(m577)* | 25°C | No carb | *daf-2(e1370)* |
| 15°C | No carb | N2 | 25°C | No carb | *daf-2(e1370)* |
| 20°C | No carb | N2 | 25°C | No carb | *daf-2(e1370)* |
| 20°C | Carb | N2 | 25°C | No carb | *daf-2(e1370)* |
| 25°C | Carb | N2 | 25°C | No carb | *daf-2(e1370)* |
| 15°C | Carb | *daf-16(mgDf50)* | 25°C | Carb | *daf-2(e1370)* |
| 20°C | Carb | *daf-16(mgDf50)* | 25°C | Carb | *daf-2(e1370)* |
| 25°C | Carb | *daf-2(m577)* | 25°C | Carb | *daf-2(e1370)* |
| 20°C | Carb | N2 | 25°C | Carb | *daf-2(e1370)* |
| 15°C | No carb | *daf-16(mgDf50)* | 15°C | No carb | *daf-2(m577)* |
| 20°C | No carb | *daf-16(mgDf50)* | 15°C | No carb | *daf-2(m577)* |
| 20°C | Carb | *daf-16(mgDf50)* | 15°C | No carb | *daf-2(m577)* |
| 15°C | No carb | *daf-16(mgDf50)* | 20°C | No carb | *daf-2(m577)* |
| 20°C | No carb | *daf-16(mgDf50)* | 20°C | No carb | *daf-2(m577)* |
| 20°C | Carb | *daf-16(mgDf50)* | 20°C | No carb | *daf-2(m577)* |
| 25°C | Carb | *daf-16(mgDf50)* | 20°C | No carb | *daf-2(m577)* |
| 15°C | No carb | N2 | 20°C | No carb | *daf-2(m577)* |
| 25°C | Carb | N2 | 20°C | No carb | *daf-2(m577)* |
| 15°C | No carb | *daf-16(mgDf50)* | 25°C | No carb | *daf-2(m577)* |
| 20°C | No carb | *daf-16(mgDf50)* | 25°C | No carb | *daf-2(m577)* |
| 25°C | No carb | *daf-16(mgDf50)* | 25°C | No carb | *daf-2(m577)* |
| 25°C | Carb | *daf-16(mgDf50)* | 25°C | No carb | *daf-2(m577)* |
| 25°C | Carb | N2 | 25°C | No carb | *daf-2(m577)* |
| 15°C | No carb | *daf-16(mgDf50)* | 15°C | No carb | N2 |
| 20°C | No carb | *daf-16(mgDf50)* | 15°C | No carb | N2 |
| 20°C | Carb | *daf-16(mgDf50)* | 15°C | No carb | N2 |
| 25°C | Carb | *daf-16(mgDf50)* | 15°C | No carb | N2 |
| 25°C | Carb | N2 | 15°C | No carb | N2 |
| 20°C | No carb | *daf-16(mgDf50)* | 20°C | No carb | N2 |
| 25°C | No carb | *daf-16(mgDf50)* | 20°C | No carb | N2 |
| 25°C | Carb | *daf-16(mgDf50)* | 20°C | No carb | N2 |
| 20°C | Carb | *daf-16(mgDf50)* | 20°C | Carb | N2 |
| 25°C | No carb | *daf-16(mgDf50)* | 25°C | No carb | N2 |

**Table S5.** | Morbidity-compressing non-S-M treatments. Temperature, antibiotic usage and genotype of non-S-M treatments that decreased relative G-span. Each treatment comprises a pair of cohorts (control and treatment cohorts), and each row contains one treatment (39 in total).

| Non-S-M treatments |  |  |  |  |  |
| --- | --- | --- | --- | --- | --- |
| Control cohort |  |  | Treatment cohort |  |  |
| Temperature | ± Carbenicillin | Genotype | Temperature | ± Carbenicillin | Genotype |
| 20°C | No carb | *daf-16(mgDf50)* | 15°C | No carb | *daf-16(mgDf50)* |
| 25°C | No carb | *daf-16(mgDf50)* | 15°C | No carb | *daf-16(mgDf50)* |
| 25°C | Carb | *daf-16(mgDf50)* | 15°C | No carb | *daf-16(mgDf50)* |
| 20°C | Carb | *daf-16(mgDf50)* | 15°C | Carb | *daf-16(mgDf50)* |
| 25°C | Carb | *daf-16(mgDf50)* | 15°C | Carb | *daf-16(mgDf50)* |
| 25°C | No carb | *daf-16(mgDf50)* | 20°C | No carb | *daf-16(mgDf50)* |
| 25°C | Carb | *daf-16(mgDf50)* | 20°C | No carb | *daf-16(mgDf50)* |
| 25°C | No carb | *daf-16(mgDf50)* | 15°C | No carb | *daf-2(e1368)* |
| 25°C | Carb | *daf-16(mgDf50)* | 15°C | No carb | *daf-2(e1368)* |
| 15°C | No carb | N2 | 15°C | No carb | *daf-2(e1368)* |
| 20°C | Carb | N2 | 15°C | No carb | *daf-2(e1368)* |
| 25°C | Carb | N2 | 15°C | No carb | *daf-2(e1368)* |
| 25°C | No carb | *daf-16(mgDf50)* | 20°C | No carb | *daf-2(e1368)* |
| 20°C | No carb | *daf-2(m577)* | 20°C | No carb | *daf-2(e1368)* |
| 20°C | No carb | N2 | 20°C | No carb | *daf-2(e1368)* |
| 25°C | No carb | N2 | 20°C | No carb | *daf-2(e1368)* |
| 20°C | Carb | *daf-16(mgDf50)* | 20°C | Carb | *daf-2(e1368)* |
| 25°C | Carb | *daf-16(mgDf50)* | 20°C | Carb | *daf-2(e1368)* |
| 20°C | Carb | N2 | 20°C | Carb | *daf-2(e1368)* |
| 25°C | No carb | N2 | 25°C | No carb | *daf-2(e1368)* |
| 25°C | No carb | *daf-16(mgDf50)* | 25°C | Carb | *daf-2(e1368)* |
| 25°C | Carb | *daf-16(mgDf50)* | 25°C | Carb | *daf-2(e1368)* |
| 25°C | Carb | N2 | 25°C | Carb | *daf-2(e1368)* |
| 25°C | Carb | *daf-16(mgDf50)* | 20°C | No carb | *daf-2(e1370)* |
| 15°C | Carb | N2 | 20°C | No carb | *daf-2(e1370)* |
| 25°C | No carb | *daf-2(m577)* | 25°C | No carb | *daf-2(e1370)* |
| 25°C | Carb | *daf-16(mgDf50)* | 25°C | Carb | *daf-2(e1370)* |
| 15°C | Carb | N2 | 25°C | Carb | *daf-2(e1370)* |
| 25°C | No carb | *daf-16(mgDf50)* | 15°C | No carb | *daf-2(m577)* |
| 25°C | Carb | *daf-16(mgDf50)* | 15°C | No carb | *daf-2(m577)* |
| 15°C | No carb | N2 | 15°C | No carb | *daf-2(m577)* |
| 20°C | Carb | N2 | 15°C | No carb | *daf-2(m577)* |
| 25°C | Carb | N2 | 15°C | No carb | *daf-2(m577)* |
| 25°C | No carb | *daf-16(mgDf50)* | 20°C | No carb | *daf-2(m577)* |
| 20°C | Carb | *daf-16(mgDf50)* | 25°C | Carb | *daf-2(m577)* |
| 25°C | Carb | *daf-16(mgDf50)* | 25°C | Carb | *daf-2(m577)* |
| 25°C | No carb | *daf-16(mgDf50)* | 15°C | No carb | N2 |
| 25°C | Carb | *daf-16(mgDf50)* | 20°C | Carb | N2 |
| 25°C | Carb | *daf-16(mgDf50)* | 25°C | Carb | N2 |

**Table S6.**
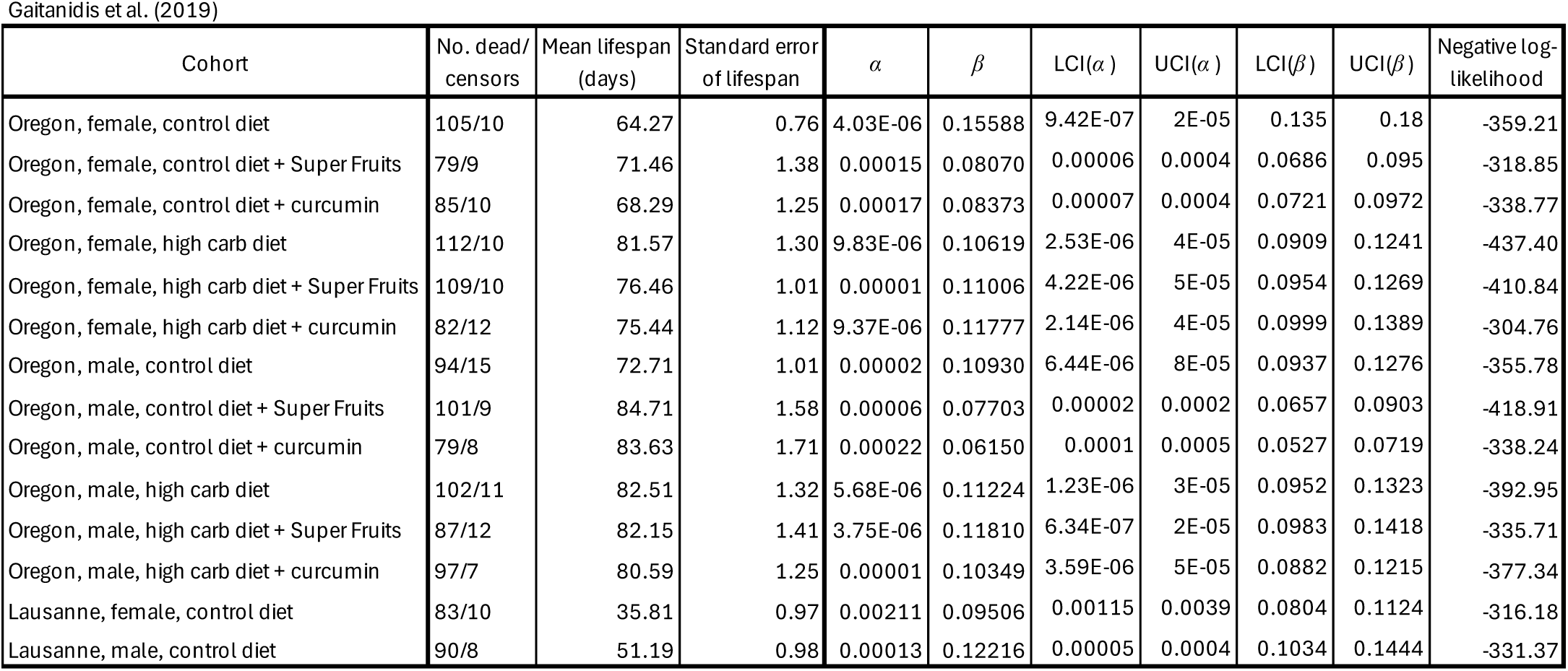
| Lifespan and Gompertz statistics for *Drosophila* cohorts in Gaitanidis et al. (**2019**). Mean lifespan and its standard errors were obtained by Kaplan-Meier survival analysis in JMP, and Gompertz parameters and their associated statistics (LCI: lower 95% confidence interval; UCI: upper 95% confidence interval) obtained by maximum likelihood estimation in WinModest.

**Table S7.** | Lifespan and Gompertz statistics for *Drosophila* cohorts in Curtsinger. (**2015**). Mean lifespan and its standard errors were obtained by Kaplan-Meier survival analysis in JMP, and Gompertz parameters and their associated statistics (LCI: lower 95% confidence interval; UCI: upper 95% confidence interval) obtained by maximum likelihood estimation in WinModest. *These data were extracted from event history graphs in Curtsinger (2015), producing 100 pseudo-individuals per cohort (see Methods).

| Cohort (strain) | No. dead/censors | Mean lifespan (days) | Standard error of lifespan | $\alpha$ | $\beta$ | LCI($\alpha$) | UCI($\alpha$) | LCI($\beta$) | UCI($\beta$) | Negative log-likelihood |
| --- | --- | --- | --- | --- | --- | --- | --- | --- | --- | --- |
| AUS | 100/0* | 38.12 | 0.87 | 0.00162 | 0.09677 | 0.00091 | 0.00288 | 0.08360 | 0.11202 | -373.80 |
| CO1 | 100/0* | 40.45 | 1.13 | 0.00140 | 0.09176 | 0.00074 | 0.00267 | 0.07786 | 0.10813 | -385.86 |
| CO2 | 100/0* | 44.16 | 1.20 | 0.00120 | 0.08589 | 0.00063 | 0.00231 | 0.07288 | 0.10121 | -393.09 |
| CO3 | 100/0* | 41.78 | 1.12 | 0.00156 | 0.08506 | 0.00085 | 0.00288 | 0.07230 | 0.10007 | -390.66 |
| LB | 100/0* | 33.28 | 1.36 | 0.00712 | 0.05564 | 0.00459 | 0.01104 | 0.04504 | 0.06874 | -409.33 |
| RI7 | 100/0* | 48.89 | 1.44 | 0.00141 | 0.07074 | 0.00076 | 0.00261 | 0.05981 | 0.08366 | -410.53 |
| S9 | 100/0* | 32.52 | 0.89 | 0.00276 | 0.09742 | 0.00159 | 0.00480 | 0.08280 | 0.11461 | -372.28 |
| SA | 100/0* | 37.63 | 0.99 | 0.00104 | 0.11050 | 0.00050 | 0.00213 | 0.09352 | 0.13057 | -371.32 |
| ZAM | 100/0* | 35.28 | 1.24 | 0.00464 | 0.06674 | 0.00286 | 0.00752 | 0.05537 | 0.08045 | -401.86 |

**Table S8.** | Lifespan and Gompertz statistics for mouse cohorts in Luciano et al. (2024). Mean lifespan and its standard errors were obtained by Kaplan-Meier survival analysis in JMP, and Gompertz parameters and their associated statistics (LCI: lower 95% confidence interval; UCI: upper 95% confidence interval) obtained by maximum likelihood estimation in WinModest.

| Cohort | No. dead/<br>censors | Mean lifespan<br>(days) | Standard error<br>of lifespan | $\alpha$ | $\beta$ | LCI($\alpha$) | UCI($\alpha$) | LCI($\beta$) | UCI($\beta$) | Negative log-likelihood |
| --- | --- | --- | --- | --- | --- | --- | --- | --- | --- | --- |
| Female C57BL/6J ad lib | 13/2 | 902.91 | 21.05 | 7.66E-07 | 0.01409 | 5.16E-09 | 0.0001 | 0.0095 | 0.0209 | -32.14 |
| Male C57BL/6J ad lib | 25/3 | 924.10 | 23.74 | 0.00003 | 0.00926 | 2.2E-06 | 0.0005 | 0.0068 | 0.0126 | -73.23 |
| Female J:DO ad lib | 26/0 | 1071.04 | 17.31 | 5.1E-06 | 0.00982 | 2.8E-07 | 9E-05 | 0.0075 | 0.0129 | -69.53 |
| Female J:DO 20% CR | 55/1 | 1179.71 | 19.12 | 2.00E-05 | 0.00761 | 2.68E-06 | 0.0001 | 0.0062 | 0.0093 | -165.13 |
| Female J:DO 40% CR | 73/6 | 1281.44 | 19.99 | 0.00004 | 0.00613 | 1.00E-05 | 0.0002 | 0.0052 | 0.0073 | -237.18 |
| Female J:DO 1 day/week IF | 33/1 | 1069.07 | 14.12 | 6.32E-07 | 0.01194 | 2.60E-08 | 2E-05 | 0.0093 | 0.0153 | -83.60 |
| Female J:DO 2 day/week IF | 48/3 | 1114.50 | 14.68 | 3.5E-06 | 0.00972 | 3.78E-07 | 3E-05 | 0.0079 | 0.0119 | -132.92 |

**Table S9.** | Lifespan and Gompertz statistics for mouse cohorts in Shahmirzadi et al. (2020). Mean lifespan and its standard errors were obtained by Kaplan-Meier survival analysis in JMP, and Gompertz parameters and their associated statistics (LCI: lower 95% confidence interval; UCI: upper 95% confidence interval) obtained by maximum likelihood estimation in WinModest.

| Cohort | No. dead/<br>censors | Mean lifespan<br>(days) | Standard error<br>of lifespan | $\alpha$ | $\beta$ | LCI($\alpha$) | UCI($\alpha$) | LCI($\beta$) | UCI($\beta$) | Negative log-likelihood |
| --- | --- | --- | --- | --- | --- | --- | --- | --- | --- | --- |
| Female AKG-treated | 20/0 | 898.55 | 21.06 | 9.23E-06 | 0.01112 | 3.32E-07 | 0.0003 | 0.008 | 0.0155 | -52.01 |
| Female control | 24/0 | 831.50 | 25.22 | 0.00005 | 0.00976 | 3.43E-06 | 0.0007 | 0.007 | 0.0135 | -66.87 |
| Male AKG-treated | 21/0 | 1020.38 | 22.79 | 4.97E-06 | 0.01032 | 1.59E-07 | 0.0002 | 0.0075 | 0.0143 | -56.30 |
| Male control | 24/0 | 976.79 | 20.30 | 3.00E-06 | 0.01137 | 9.26E-08 | 0.0001 | 0.0083 | 0.0155 | -62.62 |

**Table S10.** | Lifespan and Gompertz statistics for mouse cohorts in Borland et al. (2024). Mean lifespan and its standard errors were obtained by Kaplan-Meier survival analysis in JMP, and Gompertz parameters and their associated statistics (LCI: lower 95% confidence interval; UCI: upper 95% confidence interval) obtained by maximum likelihood estimation in WinModest.

| Cohort | No. dead/<br>censors | Mean lifespan<br>(days) | Standard error<br>of lifespan | $\alpha$ | $\beta$ | LCI($\alpha$) | UCI($\alpha$) | LCI($\beta$) | UCI($\beta$) | Negative log-likelihood |
| --- | --- | --- | --- | --- | --- | --- | --- | --- | --- | --- |
| WT females | 53/0 | 699.19 | 19.43 | 0.00064 | 0.00769 | 2.00E-04 | 0.0021 | 0.0062 | 0.0095 | -157.58 |
| Polr3b+/- females | 51/0 | 668.18 | 20.30 | 0.00103 | 0.00728 | 0.00035 | 0.0031 | 0.0059 | 0.009 | -153.65 |
| WT males | 50/0 | 784.42 | 20.84 | 0.00019 | 0.00847 | 0.00004 | 0.0009 | 0.0068 | 0.0106 | -146.15 |
| Polr3b+/- males | 52/0 | 715.60 | 23.44 | 0.00087 | 0.00693 | 2.80E-04 | 0.0027 | 0.0056 | 0.0086 | -160.69 |

## Notes

### Competing Interest Statement

The authors have declared no competing interest.

### Summary of Updates

Refinement of the definition of Strehler Mildvan correlations. Addition of two authors.

